# Trends and characteristics of multidrug resistant MRSA in Norway 2008-2020

**DOI:** 10.1101/2025.01.16.633344

**Authors:** Torunn Gresdal Rønning, Hege Enger, Jan Egil Afset, Christina Gabrielsen Ås

**Affiliations:** The Norwegian MRSA Reference laboratory, Department of Medical Microbiology, Clinic of Laboratory Medicine, St. Olavs Hospital, Trondheim University Hospital, Trondheim, Norway; Department of Clinical and Molecular Medicine, Norwegian University of Science and Technology, Trondheim, Norway; Department of Medical Microbiology, Clinic of Laboratory Medicine, St. Olavs Hospital, Trondheim University Hospital, Trondheim, Norway

**Keywords:** *Staphylococcus aureus*, MRSA, multidrug resistance, AMR, epidemiology, Norway

## Abstract

Infections caused by multidrug-resistant (MDR) bacteria are recognized as a critical One Health concern which poses a significant threat to public health, leading to increased morbidity and mortality across both high- and low-income countries. In this study, we investigated the epidemiology and molecular mechanisms of multidrug-resistant methicillin-resistant *Staphylococcus aureus* (MDR-MRSA) strains identified in Norway from 2008 to 2020, in order to gain a better understanding of the evolution and dissemination of multidrug resistance in *S. aureus*.

A total of 452 MDR-MRSA strains isolated from 429 individuals were analyzed from a dataset of 23,412 MRSA strains. Methods included epidemiological characterization, antimicrobial susceptibility testing (AST) and genetic analysis of a selection of strains using nanopore sequencing to identify antimicrobial resistance (AMR) genes and mutations, as well as their location on plasmids, SCC*mec* and other mobile genetic elements (MGEs).

The study revealed an overall increasing trend in MDR-MRSA strains, with healthcare-associated strains being more prevalent among MDR-MRSA compared to the overall MRSA population. Significant heterogeneity in *spa*-types and clonal complexes exhibiting multidrug resistance was observed, with high resistance rates against multiple antibiotic groups, particularly erythromycin, ciprofloxacin/norfloxacin, tetracycline, gentamicin, and clindamycin in addition to cefoxitin. The predominant MDR-MRSA clones included t1476/CC8, t127/CC1, t189/CC188 and t030, t037/CC239. A broad range of AMR genes and mutations were detected, linked to a wide variety of MGEs, highlighting the complex mechanisms of resistance development and dissemination within the MRSA population.

This study highlights the rising challenge posed by MDR-MRSA strains, and reveals the multifactorial nature of AMR in *S. aureus*, thus emphasizing the importance of continued surveillance, antibiotic stewardship and infection control measures, as well as global cooperation, in order to combat the spread of these multidrug-resistant pathogens.

**Author Summary:** In our study, we explored the landscape of multidrug-resistant methicillin-resistant *Staphylococcus aureus* (MDR-MRSA) in Norway from 2008 to 2020. This research is possible because it draws on a robust national surveillance system that has been active for over a decade, aimed at preventing the establishment of these dangerous pathogens in our healthcare facilities. While the overall incidence of MDR-MRSA was relatively low, we noticed an upward trend in the number of these resistant strains over time. This pattern, along with shifts in the molecular profiles of the strains, suggests that certain MDR-MRSA clones have become well-established and are spreading globally.

One of the most important findings was that the majority of MDR-MRSA strains were acquired abroad. This indicates that international travel and migration are significant contributors to the spread of these resistant strains, particularly from regions like Asia and Africa. This underscores the necessity for global collaboration in surveillance and antibiotic stewardship to combat the threat posed by these pathogens.

Additionally, we found that a high proportion of MDR-MRSA strains were associated with healthcare settings, primarily isolated from patients during hospital admissions. This is concerning, as it suggests that the most resistant strains are often found in hospitals, where vulnerable patients are at risk. The high antibiotic exposures in these environments likely contributes to the selection and spread of these resistant clones.

Interestingly, we discovered that many of the MDR-MRSA strains were detected in asymptomatic carriers rather than in clinical infections. This could be due to the strains being acquired abroad and subsequently identified through routine screening in healthcare settings. The overall potential implications for public health are however significant, especially since the resistance profiles of these strains can severely limit treatment options.

By utilizing advanced nanopore sequencing technology, we were able to delve deeper into the genetic elements responsible for antibiotic resistance, highlighting the extensive heterogeneity of resistance mechanisms among the MDR-MRSA strains. We found that resistance genes are primarily located on plasmids and other mobile genetic elements, which enhances their potential for spread among different strains. This complexity of resistance mechanisms and the adaptive strategies employed by MRSA highlight the ongoing battle against antibiotic resistance.

In conclusion, our study sheds light on the evolving landscape of MDR-MRSA, emphasizing the need for continued vigilance and coordinated efforts to mitigate the spread of these resistant strains.

## Introduction

*Staphylococcus aureus* colonizes the skin and mucosal surfaces of about 30 % of the human population [1]. This bacterium is however also an important human pathogen, causing a wide range of infections ranging from mild skin and soft-tissue infections to severe and invasive disease, such as endocarditis, osteomyelitis, bloodstream infection and sepsis [2].

*S. aureus* is furthermore a bacterial pathogen which has the capasity to incorporate a wide variety of mobile genetic elements (MGEs) making it able to adapt to different hosts and environments [3]. These MGEs, which include plasmids, transposons, bacteriophages and staphylococcal cassette chromosome (SCC) elements, can facilitate the horizontal transfer of genes that encode important virulence factors as well as antibiotic resistance determinants providing resistance against almost all the clinically relevant groups of antibiotics.

Plasmids play a pivotal role in horizontal gene transfer, significantly contributing to the dissemination of antimicrobial resistance (AMR) among bacteria [4]. Well-known examples in *S. aureus* include the widely disseminated *blaZ*-encoding plasmids that provide resistance to penicillins [5]. Furthermore, the aquistion of the staphylococcal cassette chromosome *mec* (SCC*mec*) carrying the *mecA* (or *mecC*) gene provides resistance to all beta-lactam antibiotics defining *S. aureus* as methicillin-resistant (MRSA) [6]. SCC*mec* can furthermore contain additional antibiotic resistance- and virulence genes contributing to the adaptability and pathogenicity of MRSA strains [7, 8]. Bacteriophages, or phages, are prevalent in the genome of most bacteria, often introducing additional genes that enhance virulence and antibiotic resistance [9]. In human-adapted *S. aureus* strains, Sa3int phages are particularly significant as they carry genes that help bacteria evade the immune system, thus increasing their virulence [9]. These examples illustrate the importance of horizontal gene transfer and MGEs in the dissemination of AMR and the evolution of bacterial pathogenicity in *S. aureus*.

Infections caused by multidrug-resistant (MDR) bacteria, including MRSA, are recognized as a critical One Health concern which poses a significant threat to public health [10]. These infections result in increased mortality and morbidity across both high- and low-income countries [11]. To better understand the mechanisms driving the spread of multidrug resistance in MRSA, this study aimed to examine the epidemiology and molecular mechanisms of multidrug-resistant MRSA strains identified in Norway in the period 2008 to 2020.

## Results

### Epidemiological characteristics of multidrug-resistant MRSA in Norway 2008-2020

A subset of 452 MDR-MRSA strains isolated from 429 persons were included in the study, from a total of 23,412 MRSA strains (1.9 %) (Table 1) in the study period from 2008 through 2020. Although the number of MDR-MRSA strains per year was low (ranging from 28 to 73) and with some fluctuations, we observed an overall increasing trend (Fig 1), except for the COVID-19 pandemic year 2020. This coincided with an overall increase in the total number of MRSA strains in Norway in the same period.

**Table 1.**
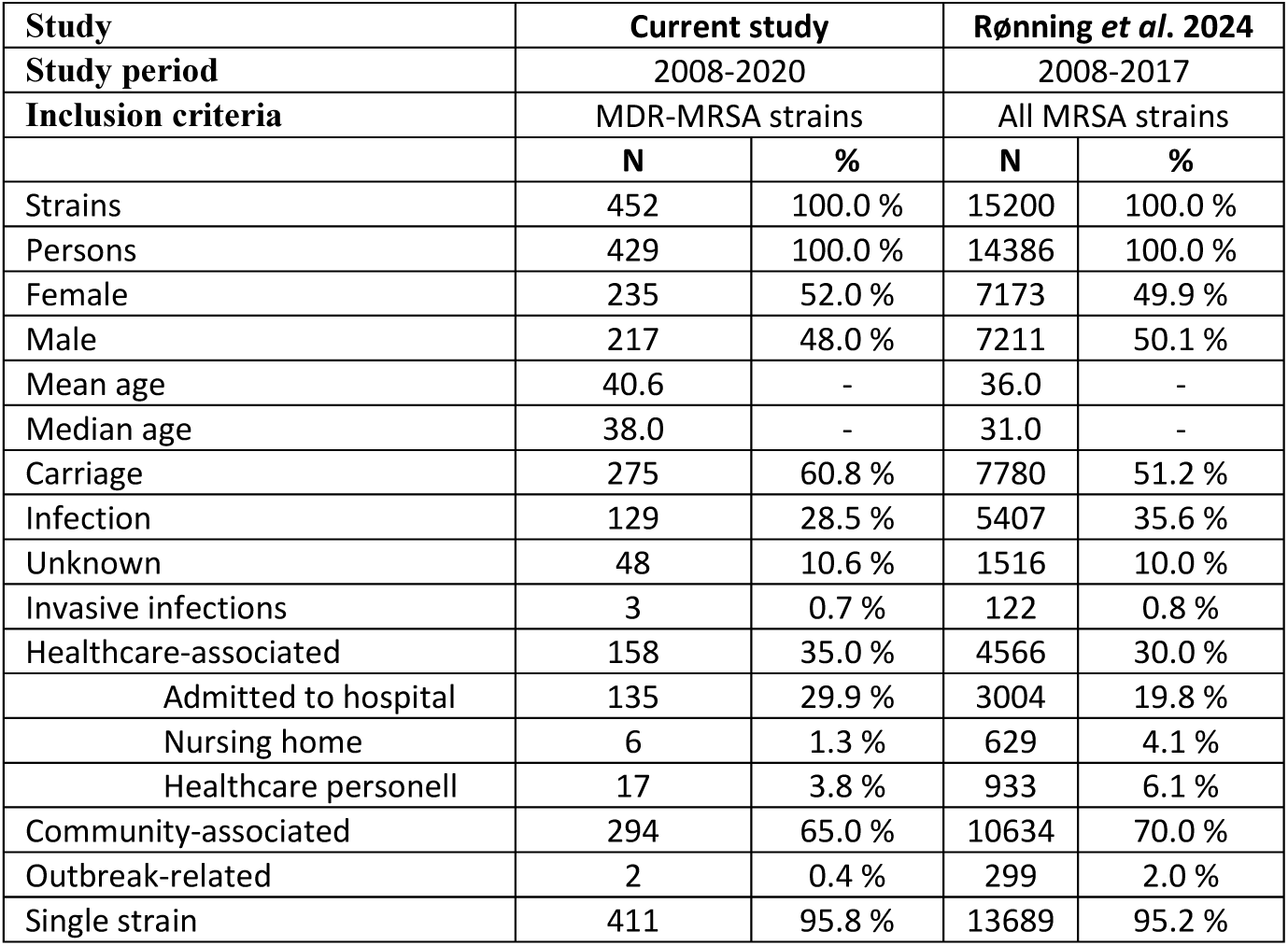
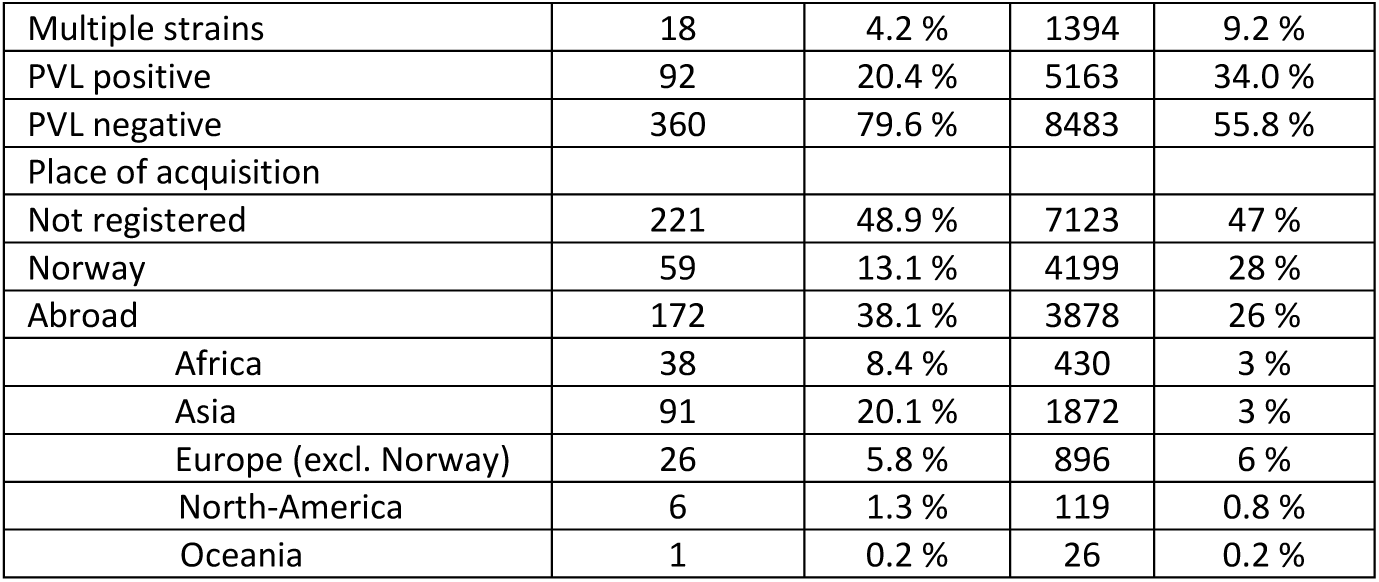
Epidemiological and molecular characteristics of the strains included in the study, compared to data from Rønning *et al*.[12] (2024).

**Fig 1.**
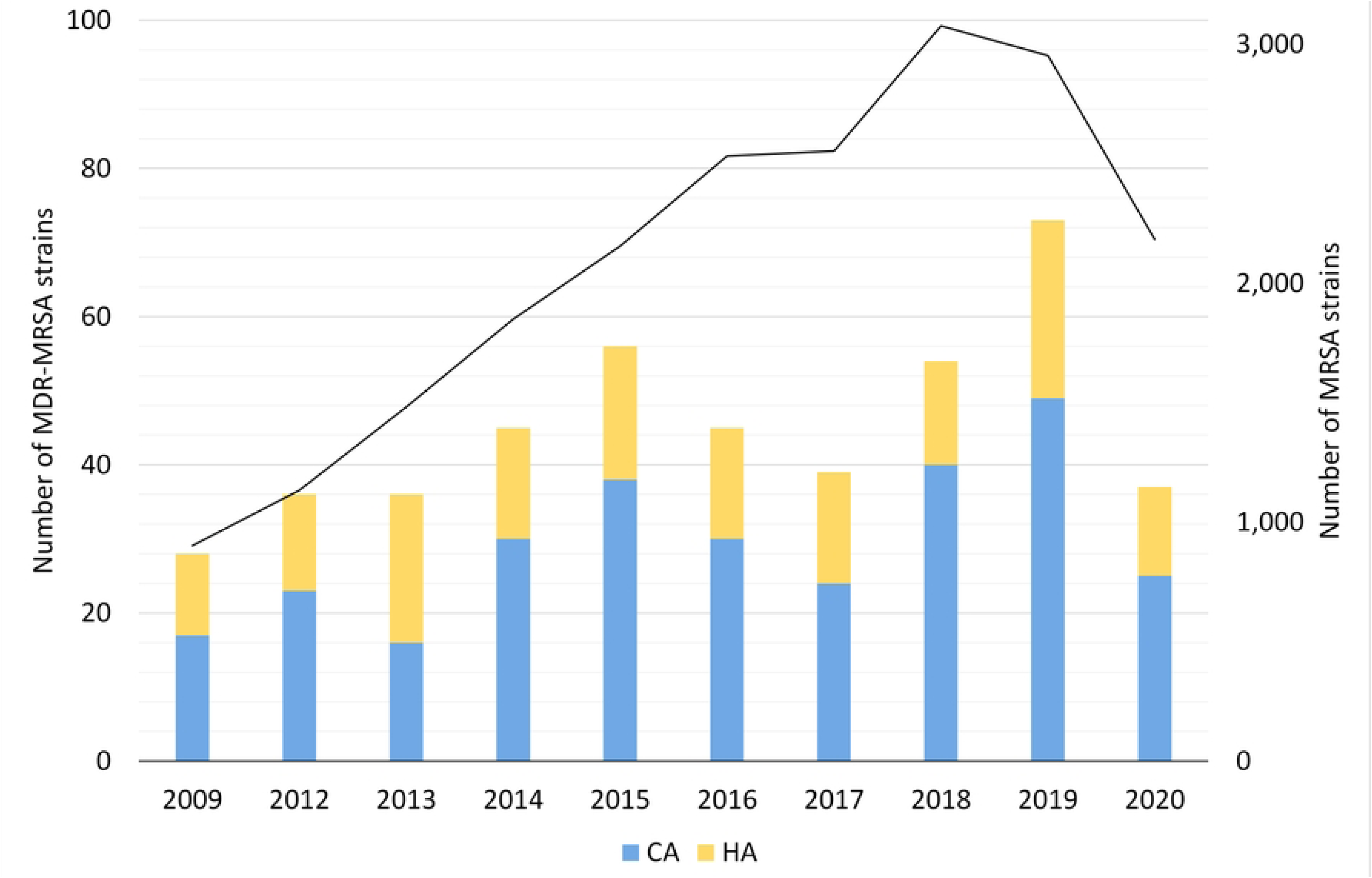
Yearly number of MDR-MRSA strains in Norway in 2008-2020. Strains are classified as Healthcare-Associated (HA) or Community-Associated (CA) as indicated by the coloured key and scale on the left axis. Total number of MRSA strains per year indicated by black line, with scale on the right axis.

In total, 275 (60.8 %) of the MDR-MRSA strains were classified as carriage strains and 129 (28.5 %) were classified as infection strains. Of the infection strains, the majority were associated with wounds (74.4 %), abcesses (14.7 %) or pus (7.8 %). In total, only three strains were from invasive infections (0.7 %). No information on sampling site was available for 48 (10.6 %) of the strains. In total, 158 strains (35.0 %) were classified as healthcare-associated (HA), and 135 strains (29.9 %) were from patients admitted to hospital. The remaining 294 strains (65.0 %) were classified as community-associated (CA). Only two of the MDR-MRSA strains (0.4 %) were registered as related to outbreaks.

According to the registered place of acquisition for the MDR-MRSA strains, 13.1 % were aquired in Norway, and 38.1 % were aquired abroad, while no information about place of acquisition was available for 48.9 % of the strains. Of the strains that were acquired abroad, the majority (20.1 %) were aquired in Asia, followed by Africa (8.4 %) and Europe (excluding Norway, 5.8 %).

The overall sex distribution of the MDR-MRSA strains was even, with 235 (52.0 %) from females, and 217 (48.0 %) from males. The mean age of persons was 40.6 years, with a median age of 38 years. More than one strain was isolated from 18 individuals (4.2 %). The strains were isolated at varying time intervals, ranging from one to six years between each collection. All strains exhibited consistent *spa*-types (or in one case clonal complex) between isolate one and isolates two or three from the same individual.

All MDR-MRSA strains in this study showed phenotypic resistance to cefoxitin and contained the *mecA*-gene, while 20.4 % (n=92) of strains contained the virulence factor and epidemiological marker Panton-Valentine leucocidin (PVL).

### Successful MDR-MRSA clones and phenotypic antimicrobial susceptibility profiles

Of the 452 MDR-MRSA strains included in the study, 361 (79.9 %) showed antibiotic resistance towards five different antibiotic groups, while 70 (15.5 %) demonstrated resistance against six antibiotic groups. Furthermore, 17 strains (3.8 %) displayed resistance against seven antibiotic groups, and four strains (0.9 %) showed antibiotic resistance against eight antibiotic groups (Fig 2).

**Fig 2.**
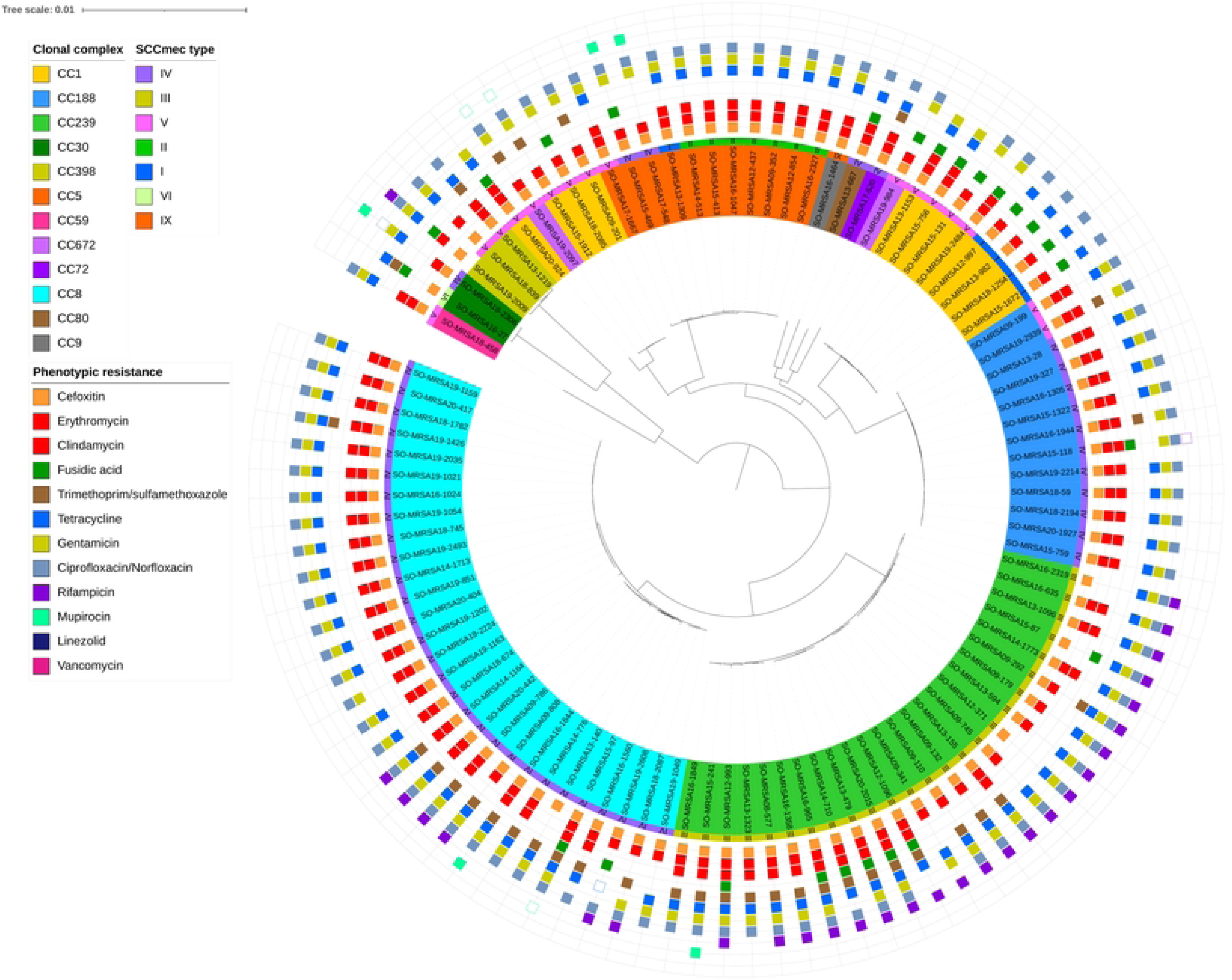
Core genome phylogeny of whole genome sequenced MDR-MRSA strains (n=101). The nodes within the tree are assigned distinct colors based on clonal complex and SCC*mec* type. Phenotypic resistance profiles based on AST are visually represented through coloured boxes (box with fill indicates resistant, while outlined box indicates intermediate resistance).

For strains showing antibiotic resistance against five or six antibiotic groups, we observed a very heterogeneous collection of *spa*-types. Strains resistant to 5 antibiotic groups belonged to more than 50 distinct *spa*-types of 22 different CCs, while strains resistant to six antibiotic groups belonged to 22 *spa*-types of 9 different CCs. Conversely, in strains showing resistance against seven or eight antibiotic groups, a very limited number of *spa*-types were observed. These included *spa*-types t008, t030, t034, t037, t064, t1476 and t451, belonging to clonal complexes 8, 239 and 398.

Overall, phenotypic antibiotic susceptibility testing revealed almost universal resistance towards erythromycin (93.1 %) and ciprofloxacin/norfloxacin (92.9 %) in addition to cefoxitine (100.0 %) (Fig 3). High levels of resistance were also observed to tetracycline (83.9 %), gentamicin (81.7 %), and clindamycin (69.3 % total). 175/452 (38.7 %) of the strains showed constitutive resistance against clindamycin, while 140 (31.0 %) showed inducible resistance against clindamycin. Moderate to low levels of resistance were observed for fusidic acid (27.8 %), trimethoprim-sulfamethoxazole (19.4 %), rifampicin (13.2 %) and mupirocin (10.0 %). No isolates were resistant towards linezolid (0.0 %) or vancomycin (0.0 %).

**Fig 3.**
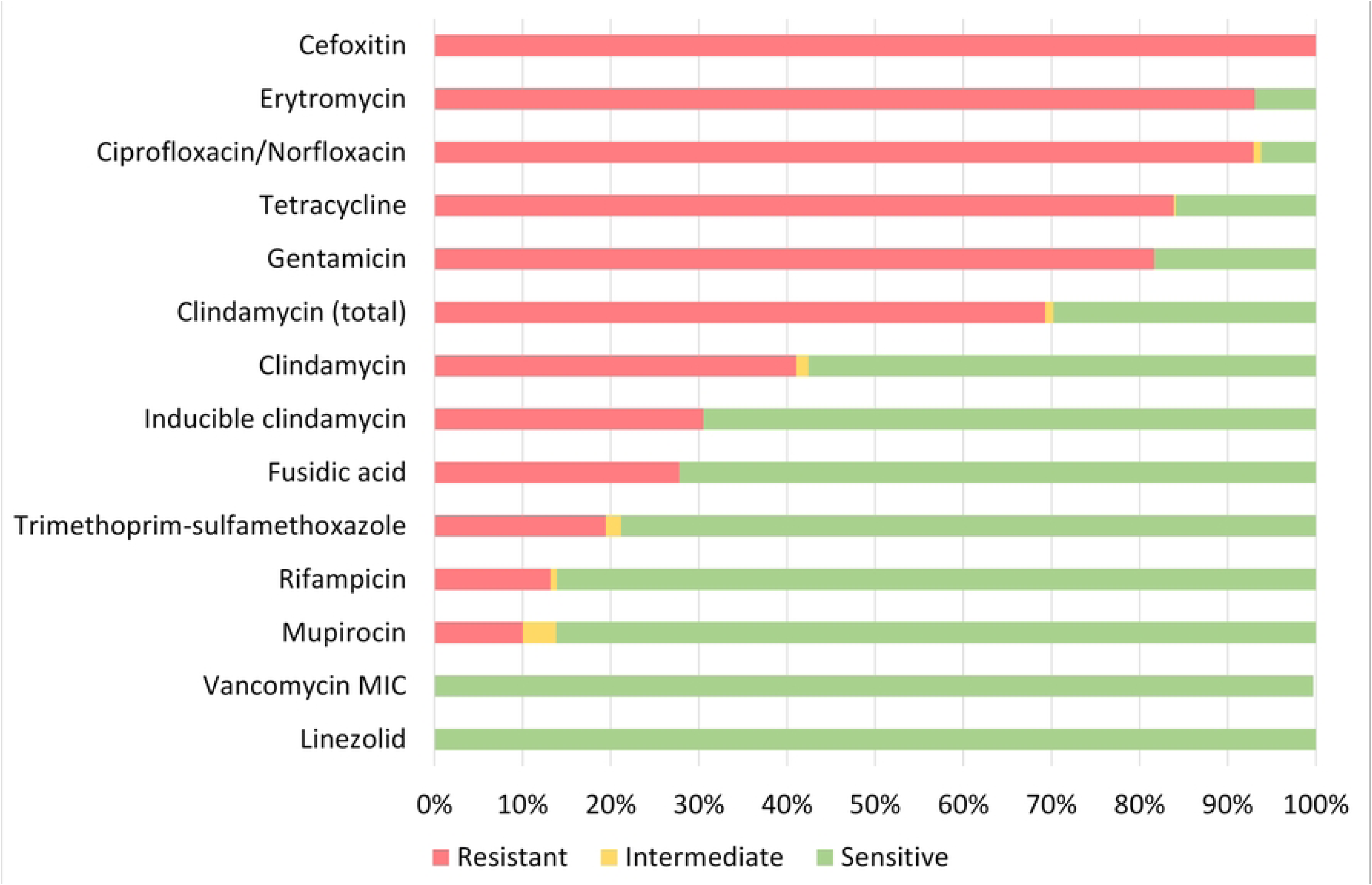
Phenotypic susceptibility of all MDR-MRSA strains categorized as resistant, intermediate or susceptible towards tested antibiotics.

Of the most successful MDR-MRSA *spa*-types over the course of the study period were t127/CC1, t189/CC188, t030, t037/CC239, and t1476/CC8 (Fig 4). Collectively, these *spa*-types accounted for 45.1 % (204/452) of the strains included in this study.

MDR-MRSA t1476/CC8 (n=63, 13.9 %) was the most successful clone in the study period (Fig 4). The number of strains which belonged to this genotype increased considerably from 2008 to 2019 (from 0 strains in 2008 to 29 strains in 2019). In this group a majority of cases were from females (65.1 %), and most of the strains were from carriage (76.2 %). Based on country of acquistion, 38.1 % of the strains were associated with countries in Sub-Saharan Africa, while 42.9 % had no record of aqcuisition (Fig 5). All the strains were resistant against beta-lactams, aminoglycosides, fluoroquinolones and tetracyclines. The majority were additionally resistant against macrolides (n=62, 98.4 %) (Table 2).

**Fig 4.**
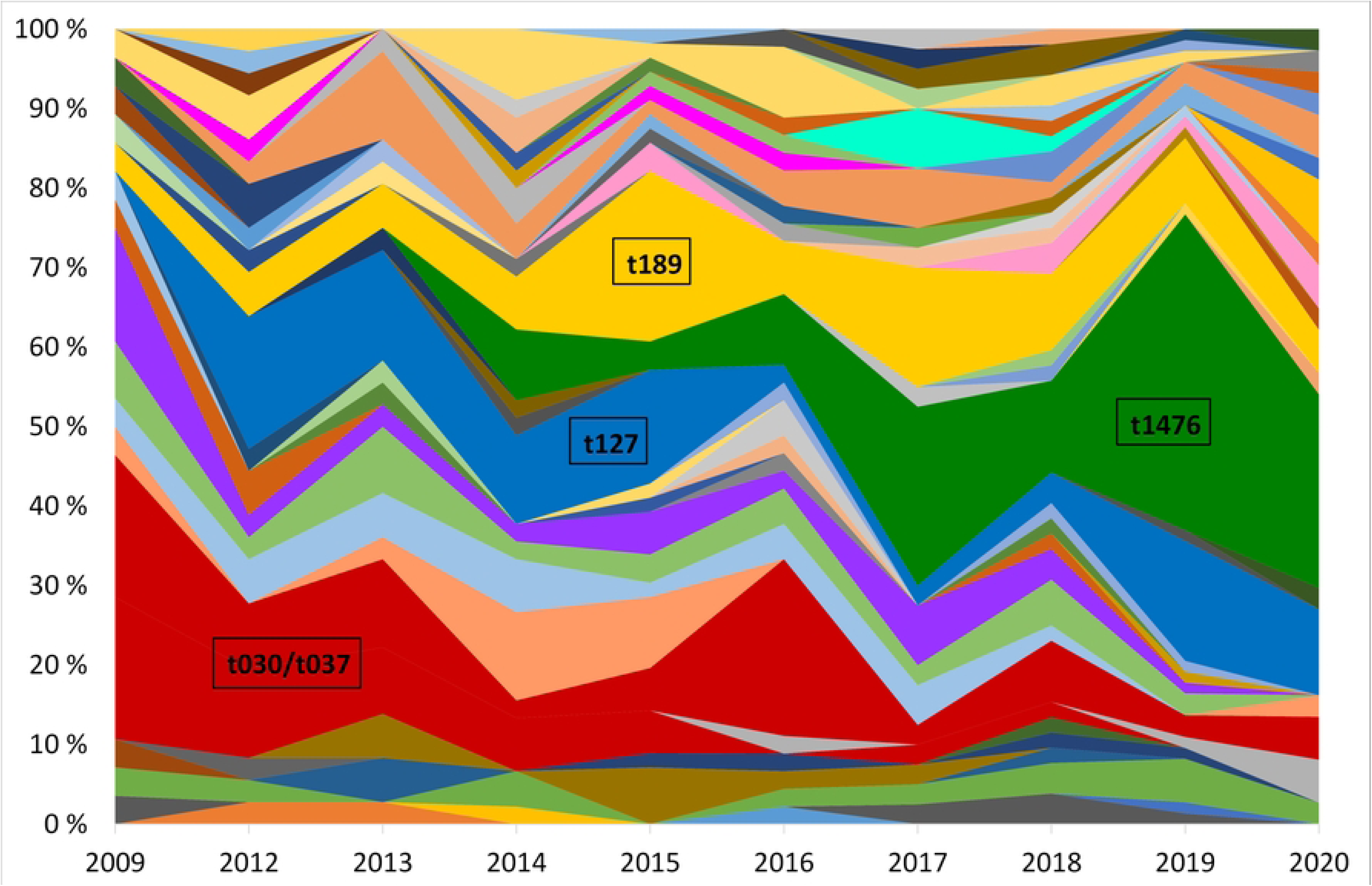
Yearly relative distribution of MDR-MRSA *spa*-types (>5 per year) in the period 2009-2020. The years 2010 and 2011 are excluded due to missing data. Major *spa*-types are highlighted.

**Fig 5.**
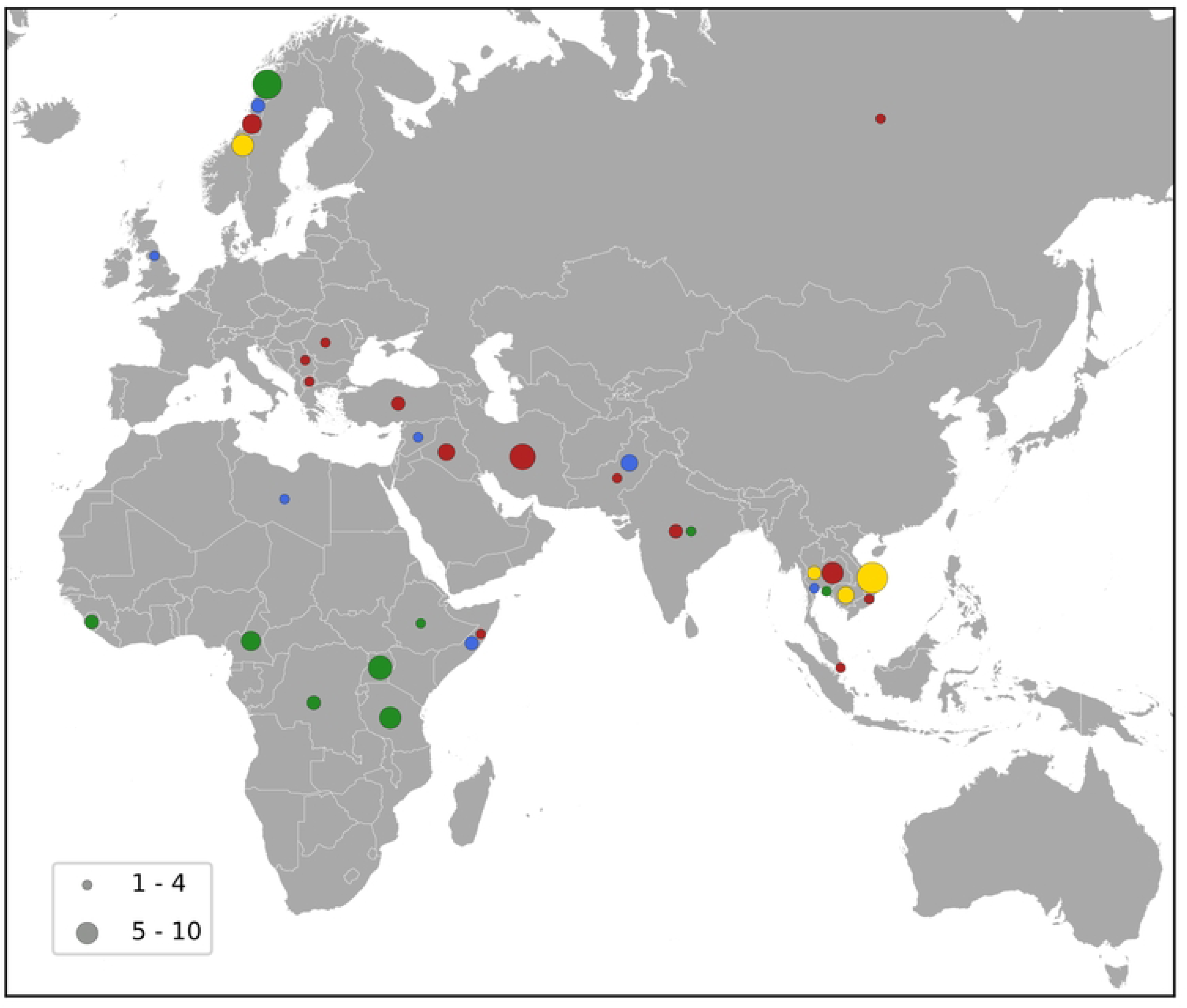
Map showing the country of aquisition for the major MDR-MRSA clones. The major clones include t1476 (green), t127 (blue), t189 (yellow) and t030/t037 (red), and the number of strains is indicated by the size of the circles as shown by the key.

**Table 2.**
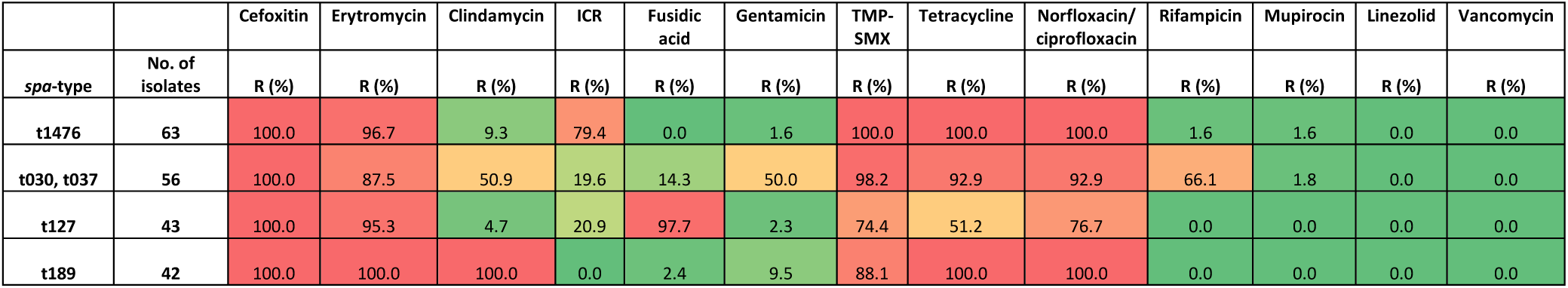
Proportion of strains resistant to the tested antibiotics, for the four most prevalent MDR-MRSA clones. Color gradient from red to green indicates high to low percentage of resistant strains accordingly. ICR, inducible clindamycin resistance; TMP-SMX, trimethoprim-sulfamethoxazole.

|  |  | Cefoxitin | Erytromycin | Clindamycin | ICR | Fusidic acid | Gentamicin | TMP-SMX | Tetracycline | Norfloxacin/ciprofloxacin | Rifampicin | Mupirocin | Linezolid | Vancomycin |
| --- | --- | --- | --- | --- | --- | --- | --- | --- | --- | --- | --- | --- | --- | --- |
| *spa*-type | No. of isolates | R (%) | R (%) | R (%) | R (%) | R (%) | R (%) | R (%) | R (%) | R (%) | R (%) | R (%) | R (%) | R (%) |
| t1476 | 63 | 100.0 | 96.7 | 9.3 | 79.4 | 0.0 | 1.6 | 100.0 | 100.0 | 100.0 | 1.6 | 1.6 | 0.0 | 0.0 |
| t030, t037 | 56 | 100.0 | 87.5 | 50.9 | 19.6 | 14.3 | 50.0 | 98.2 | 92.9 | 92.9 | 66.1 | 1.8 | 0.0 | 0.0 |
| t127 | 43 | 100.0 | 95.3 | 4.7 | 20.9 | 97.7 | 2.3 | 74.4 | 51.2 | 76.7 | 0.0 | 0.0 | 0.0 | 0.0 |
| t189 | 42 | 100.0 | 100.0 | 100.0 | 0.0 | 2.4 | 9.5 | 88.1 | 100.0 | 100.0 | 0.0 | 0.0 | 0.0 | 0.0 |

MDR-MRSA t127/CC1 was the second most frequent *spa*-type in this study, accounting for 43 out of 452 strains (9.5 %). All of these strains were resistant against five antibiotic groups (Table 2), the most common profile being resistance to beta-lactams, MLS, tetracyclines, fusidanes and fluoroquinolones (n=20, 46.5 %). The sex distribution was 48.8 % female and 51.1 % male, and the proportion of carriage (51.1 %) was similar to infections (48.8 %). For most of the cases from whom MDR-MRSA t127/CC1 strains were isolated, there was no record of place of aquisition (79.0 %). Known countries of aquisition however included European as well as Asian and African countries (Fig 5).

MDR-MRSA t189/CC188 was the third most frequent *spa*-type in this study, accounting for 42 out of 452 strains (9.3 %). These strains showed resistance against five antibiotic groups, the most common profile included resistance to beta-lactams, MLS, aminoglycosides, tetracyclines and fluoroquinolones (n=37, 88.1 %) (Table 2). The proportion of female cases was high in this group (69.0 %), and carriage (57.1 %) was more frequent than infection (40.5 %). For 35.8 % of strains, the place of acquistion were Southeast Asian countries (Fig 5).

MDR-MRSA t037/CC239 and t030/CC239 were the fourth (n=36, 8.0 %) and fifth (n=20, 4.4 %) most frequent *spa*-types in this study. These were most frequent at the start of the study period, while the number of strains declined in more recent years. These two *spa*-types belong to the same clonal complex (CC239) and share similar phenotypic antibiotic resistance patterns. A majority of strains in the two groups were resistant towards tetracyclines (98.2 %), aminoglycosides (92.9 %), MLS (87.5 %), fluoroquinolones (92.9 %), TM/S (50.0 %) and ansamycins (66.1 %) (Table 2). Notably, two strains (3.6 %), both t037, were resistant against all the antibiotics tested except for linezolid and vancomycin. In contrast to the other successful clones, there were more men (64.3 %) than women (42.9 %) in this group, and a majority of strains were from carriage (62.5 %). 42.9 % of the strains were registrered as acquired in Asian countries (Northern, Western, Southern, and South-eastern subregions) (Fig 5).

### Genotypic resistance determinants associated with different groups of antibiotics

For the sequenced MDR-MRSA strains (n=101), the presence of AMR-associated genes and mutations were predicted using the AMRFinder Plus tool and database. In total, 39 different AMR genes and 31 different AMR mutations were identified, with a median of 10 (range 7-15) different genes and 3 (range 1-10) mutations per strain. In accordance with phenotypic restistance data, the most common AMR determinants detected provided resistance against beta-lactams, MLS, fluoroqinolones, tetracyclines and aminoglycosides (Fig 2, Table 3). The specific resistance determinants associated with each group of antibiotics are more closely described in the following subsections.

**Table 3:**
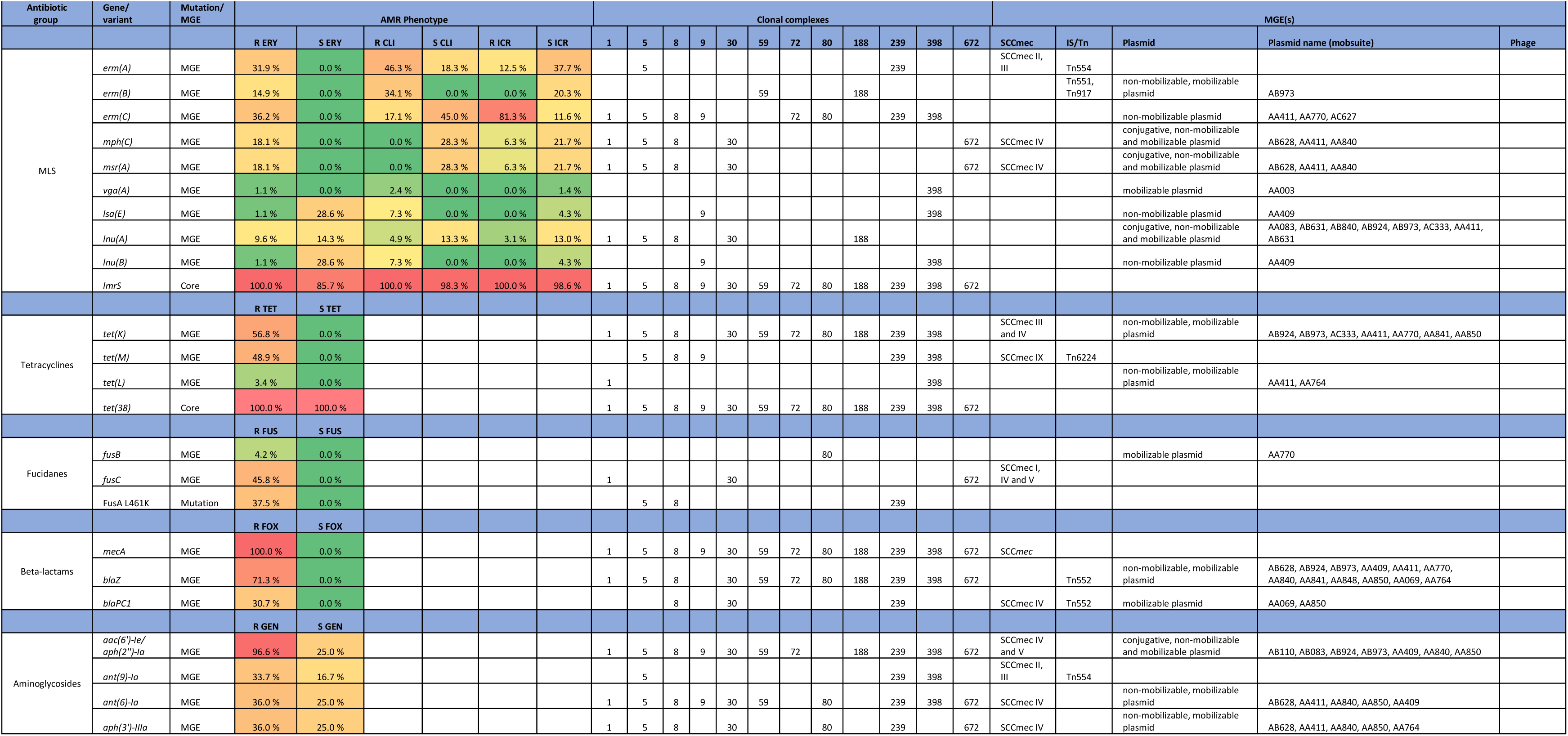
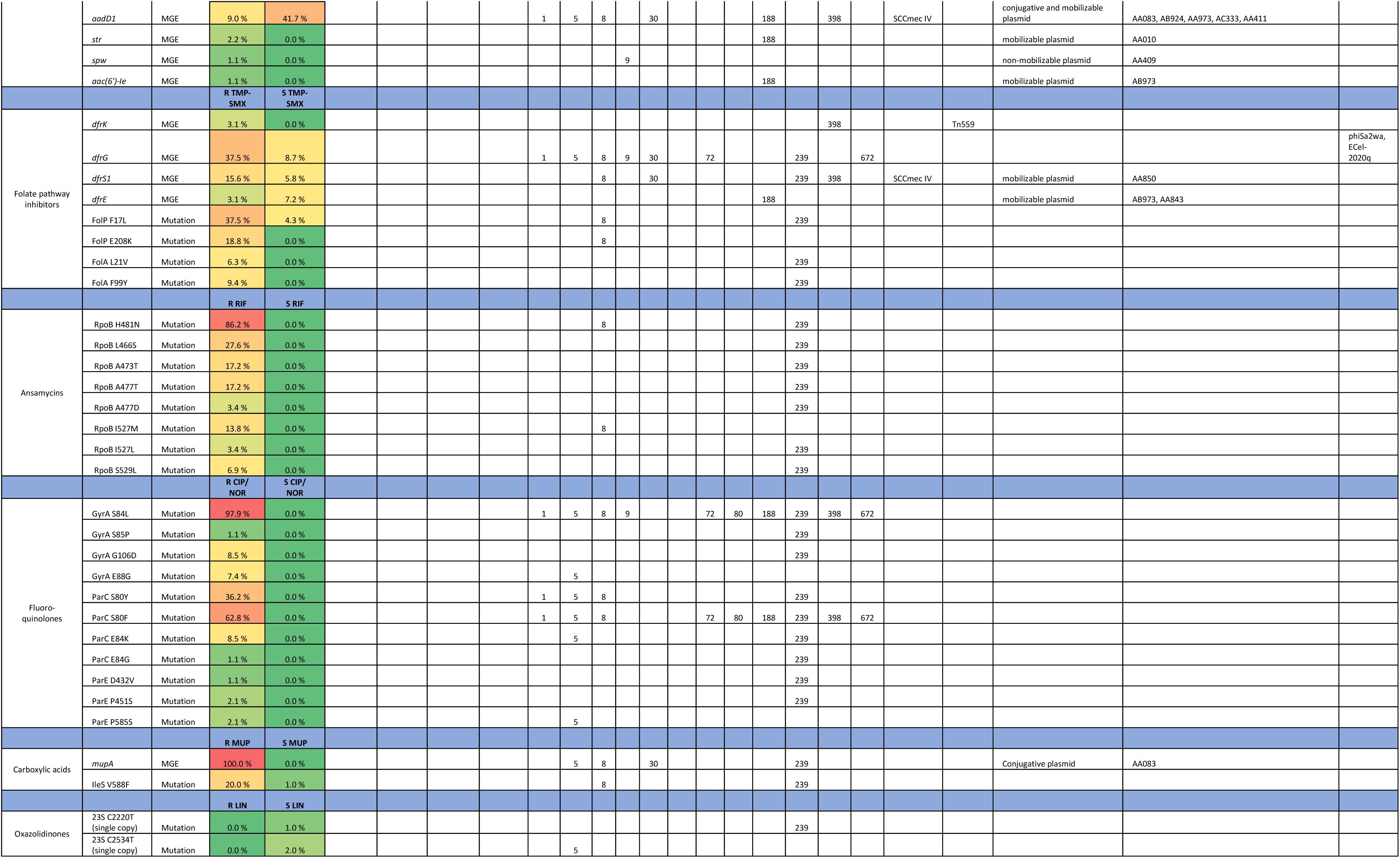
AMR traits detected in MDR-MRSA strains in Norway, and association with clonal complex and detected MGEs. AMR phenotype is given as percentage 231 of strains with the specific AMR trait where AST is interpreted as resistant (R), susceptible with increased exposure (I) or susceptible (S) towards the specific antibiotic(s) tested in each group: GEN, gentamicin; TET, tetracycline; ERY, erythromycin; CLI, clindamycin; ICR, Inducible clinidamycin resistance; FUS, fusidic acid, NOR, norfloxacin; CIP, ciprofloxacin; RIF, rifampicin; FOX, cefoxitin; TMP-SMX, Trimethoprim-sulfamethoxazole; MUP, mupirocin; LIN, linezolid.

#### Beta-lactam resistance

All MDR-MRSA strains included in this study (n=452) harboured the *mecA*-gene. In addition, we detected penicillin resistance genes in 98.0 % of the sequenced strains (Table S1). Of these, 71.3 % harboured the *blaZ*-gene, and 26.7 % contained *blaPC1*. 4 % of strains had *mecA*, *blaZ* and *blaPC1* genes, while two strains (2 %) solely harboured the *mecA*-gene.

#### Macrolide and lincosamide resistance

Almost all (95.0 %) of the sequenced MDR-MRSA strains were phenotypically resistant against erythromycin and clindamycin (Table S1). This was however linked to different genes and gene combinations. The most common gene was *erm(C)*, detected in 35.4 % of MLS-resistant strains (Table 3, Fig 6), followed by *erm(A)*, detected in 31.3 % of MLS-resistant strains. The different genetic profiles were not associated with distinct MLS phenotypic profiles, but rather included erythromycin resistance with either constitutive or inducible resistance or susceptibility to clindamycin. The *erm(B)* gene was found in 14 strains (Table 3, Table S1), and all of these had a phenotypic profile of erythromycin resistance and constitutive clindamycin resistance.

**Fig 6.**
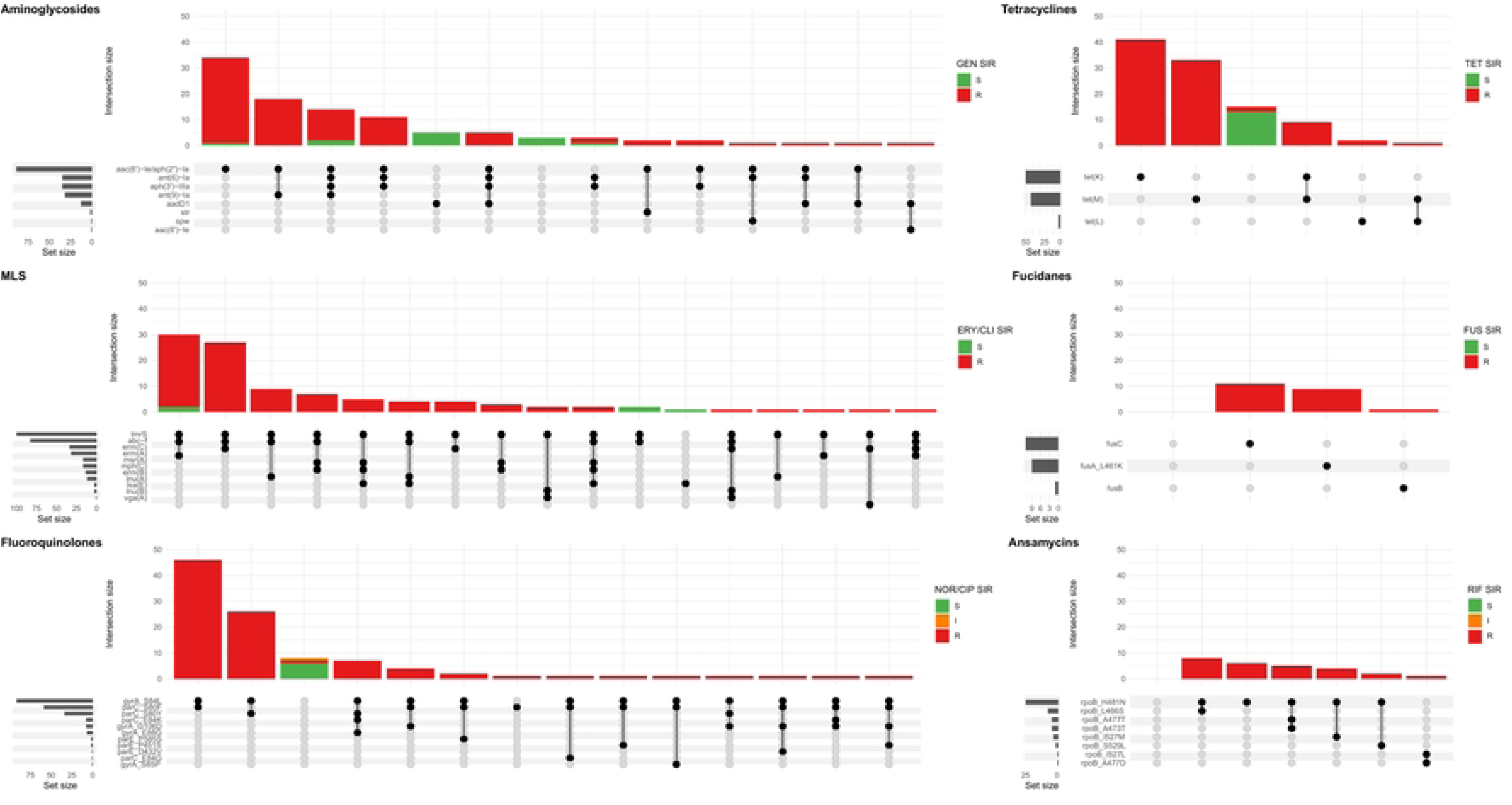
Upset plot showing all combinations AMR genes and mutations (set) associated with the major antibiotic groups in sequenced MDR-MRSA strains. The major antibiotic groups include aminoglycosides, tetracyclines, macrolide, lincosamide and streptogramin (MLS), fucidanes, fluoroquinolones and ansamycins. Strains (intersection) are colored according AST, interpreted as resistant (R), susceptible with increased exposure (I) or susceptible (S) towards the specific antibiotic(s) tested in each group: GEN, gentamicin; TET, tetracycline; ERY, erythromycin; CLI, clindamycin; FUS, fusidic acid, NOR, norfloxacin; CIP, ciprofloxacin; RIF, rifampicin.

The combination of *mph(C)* and *mrs(A)* was present in the erythromycin resistant and inducible clindamycin resistant-profile, and the only erythromycin resistant profile (Fig 6). The rarest genes associated with MLS resistance were *vga(A)* and the combination of *lsa(E)* and *lnu(B),* with the phenotypic profile erythromycin susceptible and constitutive clindamycin resistance. The gene *lmrS* was present in both erythromycin resistant and susceptible strains, and did not appear to provide phenotypic resistance to erythromycin at levels sufficient for detection by the methods used in this study.

#### Ciprofloxacin/norfloxacin resistance

Of the sequenced MDR-MRSA strains, 94 (93.1 %) exhibited phenotypic resistance to ciprofloxacin or norfloxacin. The most frequent mutations found associated with quinolone resistance were GyrA S84L (91 strains, 90.1 %), ParC S80F (58 strains, 57.4 %) and ParC S80Y (34 strains, 33.7 %) (Table 3, Table S1). Combinations of quinolone-conferring mutations were found in 92 strains (91.1 %). The most frequent combinations of mutations were GyrA S84L and ParC S80F (found in 46 strains, 45.5 %) (Fig 6) and GyrA S84L and ParC S80Y (found in 26 strains, 25.7 %). Mutations in the chromosomal *gyrA, parC* and *parE* genes were found in all quinolone-resistant strains, among multiple *spa*-types of different clonal complexes. This thus appears to be a quite common adaptation in the general MDR-MRSA population to acquire quinolone resistance.

#### Tetracycline resistance

88 (87.1 %) of sequenced MDR-MRSA strains exhibited phenotypic resistance to tetracycline, having either the *tet(M)* (42.6 %), *tet(K)* (49.5 %) or in a few cases the *tet(L)* (3.0 %) genes in different combinations (Table 3, Table S1, Fig 6). In two strains we did not detect any gene(s) likely causing tetracycline resistance. Previously described chromosomal mutations associated with tetracycline resistance in MepA (N369Y), RpsJ (Y58D) or 16S rDNA, were not identified in any of the strains. The *tet(38)-*gene was detected in all the sequenced strains in this study, but did not appear to cause phenotypic tetracycline resistance at levels sufficient for detection by the methods used in this study (Table 3).

#### Gentamicin resistance

Of the sequenced MDR-MRSA strains, 89 (88.1 %) exhibited phenotypic resistance to gentamicin. The most common gene encoding aminoglycoside resistance were *aac(6’)-Ie/aph(2’’)-Ia*, found in 86 of the 89 (96.9 %) (Table 3, Table S1). 35 strains (34.7 %) had the *aph(3’)-IIIa*-gene encoding amikacin/kanamycin resistance, 35 strains (34.7 %) had *ant(6)-Ia*-gene encoding streptomycin resistance and 32 (31.7 %) had *ant(9)-Ia*-gene encoding spectinomycin resistance. 13 strains (12.9 %) had *aadD1*-gene encoding kanamycin/tobramycin resistance. Single strains had the *aac(6’)-Ie*-gene encoding amikacin/kanamycin/tobramycin-resistance and the aminoglycoside *spw*-gene, and three strains had the *str*-gene encoding streptomycin. The most frequently found combination of aminoglycoside-genes was *aac(6’)-Ie/aph(2’’)-Ia* together with the *ant(9)-Ia*-gene, found in 18 strains (17.8 %) (Fig 6).

#### Trimethoprim-sulfamethoxazole resistance

Overall, 34 (33.7 %) of sequenced MDR-MRSA strains exhibited phenotypic resistance to TMP-SMX. As phenotypic testing is performed with the TMP-SMX combination drug, and resistance to both components separately is required for resistance, the detection and interpretation of genes and mutations contributing to TMP-SMX resistance can be complex. While we observed full concordance between phenotypic susceptibility and lack of previously described genes/mutations encoding TMP-SMX resistance, there was only a 23/34 (67.6 %) concordance between phenotypic resistance and having a combination of genes/mutations previously reported to confer phenotypic TMP-SMX resistance (Table 3, Table S1). In the concordant cases cases, the *drfG, dfrS1, dfrK* or *dfrE* genes encoding trimetoprim resistance were detected, either in a single, two or three copies, toghether with the F17L and E208K-mutations in FolP providing sulfametoxazole resistance (Table S1).

#### Rifampicin resistance

Overall, 26 (25.7 %) of sequenced MDR-MRSA strains exhibited phenotypic resistance to rifampicin. Chromosomal mutations in *rpoB* conferring rifampicin resistance described by Guerillot *et al.* were found in 26 of the sequenced strains [13]. The most common mutations were RpoB H481N alone or in combination (Table 3, Fig 6, Table S1). The chromosomal *rpoB*-mutations conferring rifampicin resistance seemed to be clonal, and were found only in strains belonging to CC8 and CC239, which included the most resistant strains in this study with resistance towards seven or eight antibiotic groups. The H481N mutation has furthermore been reported to promote the emergence of a subpopulation of small colony variants with reduced susceptibility to vancomycin and daptomycin. Although not investigated here, this raises an additional concern regarding the use of rifampicin treatment and the possible effect on emergence of MDR-MRSA strains.

#### Fusidic acid resistance

Of the sequenced MDR-MRSA strains, 25 (24.8 %) exhibited phenotypic resistance to fusidic acid. The most common gene that was associated with fusidic acid resistance was *fusC*, found in 11 strains (Table 3, Table S1). The *fusB-*gene was present in one strain (t044, CC80). The *fusA* L461K mutation was found in nine strains, were *spa*-type t037 was the most frequent *spa*-type. For three strains, we did not detect any genetic determinants likely causing phenotypic fusidic acid resistance. These strains however had disk diffusion inhibition zones close to the susceptibility breakpoint (S>24 mm). There was 100 % concordance for phenotypic susceptibility to fusidic acid and having no genetic determinants detected.

#### Mupirocin resistance

Of the sequenced MDR-MRSA strains, five (5.0 %) exhibited phenotypic resistance to mupirocin (Table 3, Table S1). All of these strains contained the *mupA*-gene. One strain had two copies of the gene, and one strain furthermore had the chromosomal *IleS* V588F mutation in addition to the *mupA*-gene. The V588F IleS-mutation was however also detected in one mupirocin susceptible strain.

#### Linezolid resistance

No phenotypic resistance to linezolid was observed in this study, and no genes associated with linezolid resistance (*cfr, optrA, poxtA*) were detected in the sequenced MDR-MRSA strains. Linezolid resistance may also be caused by mutations in copies of the 23S rDNA genes (7). We identified two mutations associated with linezolid resistance in three MDR-MRSA strains; C2192T (n=1) and C2534T (n=2) [14, 15] (Table 3). However, neither of the strains demonstrated phenotypic resistance when exposed to linezolid, as all had mutations in only a single copy of the gene.

### Mobile genetic elements associated with antibiotic resistance determinants

For the sequenced MDR-MRSA strains (n=101), the presence of AMR genes within specific types of MGEs were investigated. The specific resistance determinants associated with each type of MGE are more closely described in the following subsections.

#### Plasmids

In the present study, a total of 191 plasmids were identified, in 83.2 % of the sequenced MDR-MRSA strains. The number of plasmids per strain varied from 1 to 6, with a median of 2 plasmids per strain, and a median size of 10,318 bp. The mobilizable group of plasmids included both the largest and some of the smallest plasmids, ranging in size from 1,299-98,879 bp (Fig S1). A majority of the plasmids were previously described in *S. aureus* (86.4 %) or other Staphylococci (10.5 %). However, plasmids previously described in *Escherichia coli* (2.1 %), *Lactobacillus pentosus* (0.5 %) and *Salmonella enterica* 0.5 %) were also detected.

The plasmids were clustered into 29 sub communities using Pling, while two of the nodes were excluded due to likely being partial plasmids or transposons. Among these sub communities, 15 comprised of multiple plasmids while 17 were singletons. The largest sub community included 117 plasmids, indicating that a majority of the plasmids in this study were related. The largest sub community included plasmids from strains of 25 different *spa*-types and 11 different sequence types.

These plasmids encoded various combinations of 21 different AMR-genes, covering resistance against beta-lactams, MLS, aminoglycosides, trimethoprim, tetracyclines, fosfomycins and fusidanes. The second largest sub community included 11 plasmids, from strains with five different *spa*-types, and four different sequence types. These plasmids contained combinations of five different AMR-genes, covering resistance against beta-lactams, aminoglycosides, lincosamide and mupirocin.

The majority of plasmids were convergent plasmids (48.4 %), carrying combinations of AMR-, stress and virulence-genes (e.g. AA411) (Table S2). Some of the detected plasmids were however strictly AMR-(32.3 %), stress-(12.9 %) or virulence-plasmids (6.5 %), carrying genes of only one specific category (e.g. AC627, AA851 and AA379, respectively). In total, 112 plasmids (58.6 %) carried one or several AMR genes, with a median of 4 AMR genes per plasmid (ranging from 1 to 14). The AMR genes most commonly found on plasmids were *blaZ* (n=45), *aac(6’)-Ie/aph(2’’)-Ia* (n=30), *erm(C)* (n=26) and *tet(K)* (n=24). The *blaZ* gene was found on 15 different plasmids, the *aac(6’)-Ie/aph(2’’)-Ia* on nine different plasmids, the *erm(C)* gene on three different plasmids and the *tet(K)* gene on eight different plasmids (Fig 7). Other genes were only found on specific plasmids; e.g. *mup(A)* was only found in the conjugative plasmid AA083. The most common AMR plasmids identified in the MDR-MRSA strains were the non-mobilizable plasmid AC627 (n=20), and the mobilizable plasmids AA411, AB973 and AC333 (n=13). Conjugative AMR plasmids included the AA083 (n=4) and the AB110 (n=3) plasmids.

**Fig 7.**
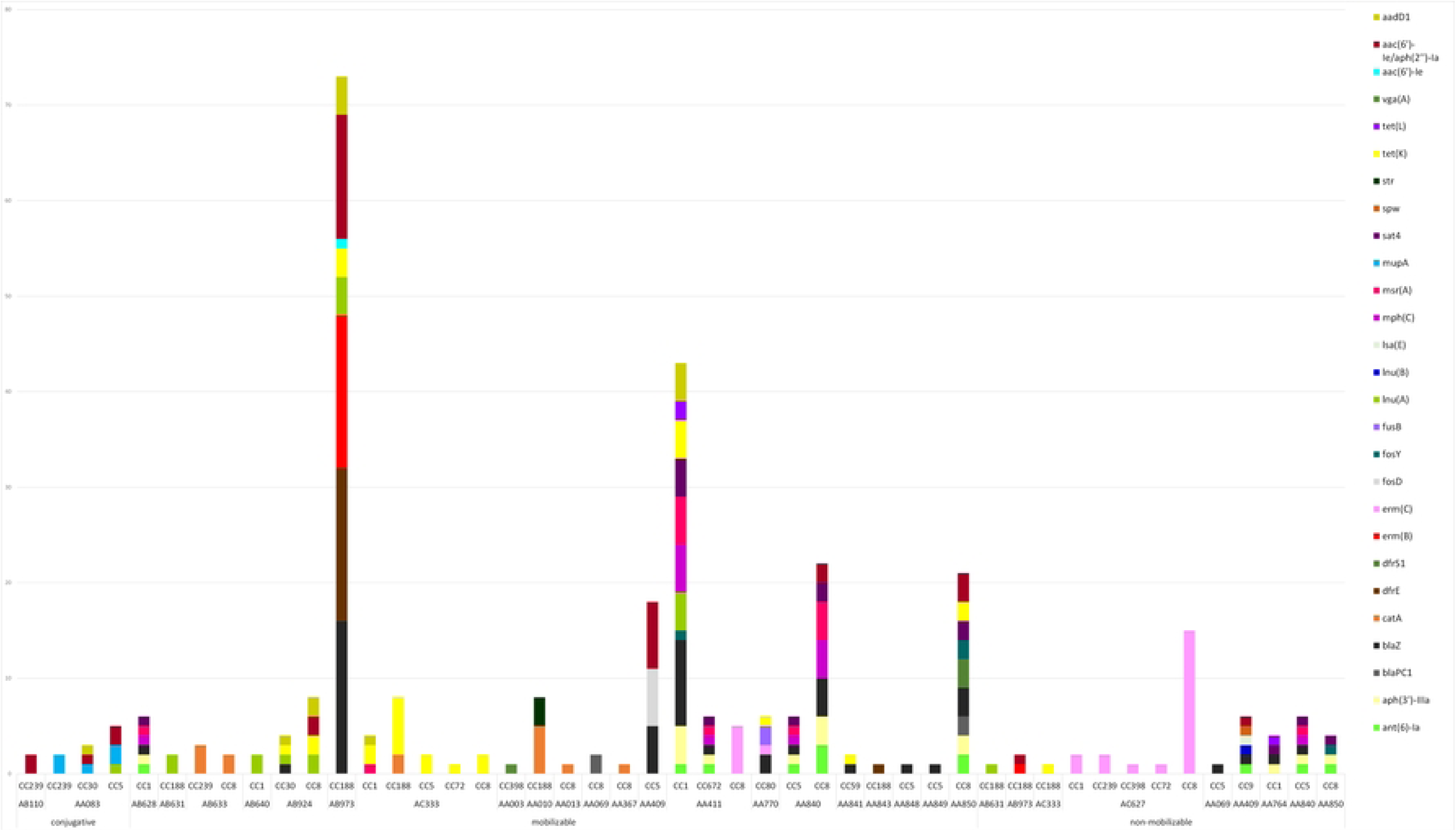
Plasmid type, predicted mobility and AMR genes found within plasmids in sequenced MDR-MRSA strains from Norway.

The small non-mobilizable plasmid AC627 (Fig 7, Table S2), which only holds the *erm(C)*-gene, was found in multiple strains belonging to CC1 (t127), CC239 (t037 and t632), CC398 (t011), CC72 (t3092) and CC8 (t064, t1476, t1952 and t451). For seven strains where we initially detected no genetic cause of macrolide resistance, it was upon further inspection detected that this plasmid was present, but had not been assembled correctly. Previous studies have also reported challenges in the detection of small plasmids using long read assemblers [16]. Consequently, this is an important aspect to consider for analysis of small plasmids when employing this methodological approach.

Most of the detected plasmids (76.0 %) were found in a single clonal complex (Fig 7), and thus appeared to be relatively stably maintained within a specific genetic background while not being disseminated to other clones to any large extent. Consequently, these plasmids likely have diminished capacity for disseminating AMR traits across the more widely distributed MRSA clones. For instance, the medium sized mobilizable plamid AB973, holding the AMR genes *erm(B), blaZ, blaR1, blaI, dfrE, lnu(A), tet(K), aac(6’)-Ie/aph(2’’)-Ia* and *aadD1,* was only found in strains belonging to CC188 (Fig 7, Table S2). A few groups of AMR-plasmids (18.8 %) however appeared to be spreading more successfully to diverse MRSA backgrounds. The medium sized conjugative plasmid AA083, holding *mup(A), lnu(A), qacC, aac(6’)-Ie/aph(2’’)-Ia* and *aadD1*, was found in CC239 (t037), CC30 (t665) as well as CC5 (t067 and t9408). Furthermore, the small plasmids AC333 and AC627, the small to medium-sized plasmids AA411, AB924 and AA840, were found in multiple clonal backgrounds, and thus demonstrate the highest potential for horizontal transfer between different MRSA clones.

#### Staphylococcal chromosome cassette mec (SCCmec)

SCC*mec* was detected in all but one (99.0 %) of the sequenced MDR-MRSA strains. The lengths of SCC*mec* chromosome cassettes were quite uniform, with a median length of 45,587 bp. The smallest and largest SCC*mec* elements detected were both SCC*mec* type IV, of length 24,060 bp and 83,838 bp, respectively. The most prevalent SCC*mec* types identified in the sequenced strains were type IV (n=47, 46.5 %) followed by type III (n=25, 24.8 %). Both SCC*mec* types contained a wide variety of AMR genes (Fig 8). Type IV was typically detected in strains belonging to CC8 and CC188. The most commonly detected AMR genes in this SCC*mec*, in addition to *mecA*, were *aac(6’)-Ie/aph(2’’)-Ia*, *drfS1* (n=17), and *tet(K)* (n=16). SCC*mec* type III was only detected in strains belonging to CC239. The most predominant AMR genes found within this type, aside from *mecA,* were *ant(9)-Ia* and *erm(A)* (n=14).

**Fig 8.**
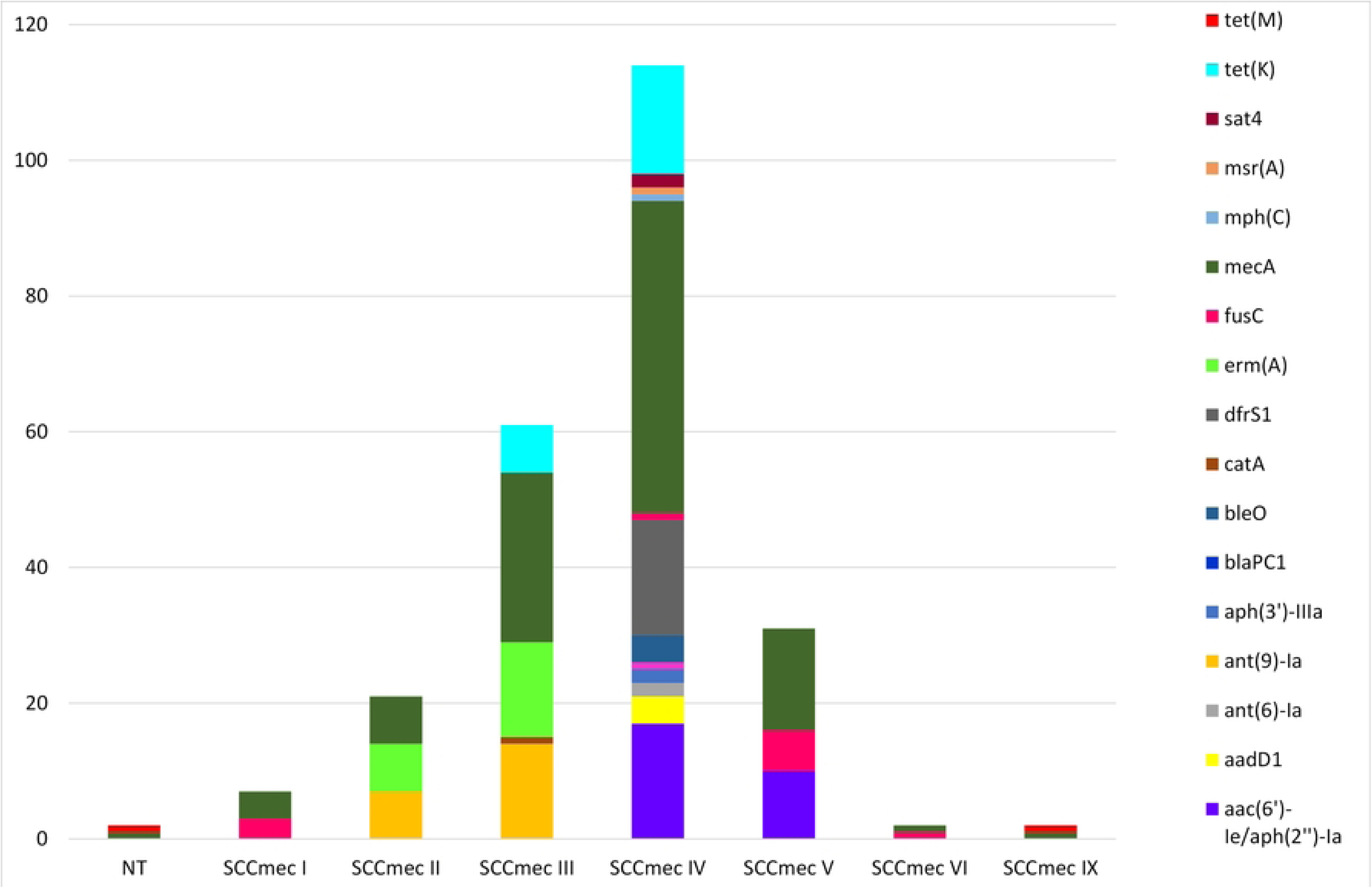
AMR genes identified in SCC*mec* elements in sequenced MDR-MRSA strains from Norway. NT, non-typeable.

#### Prophages

A total of 256 prophages harboring virulence and/or AMR-genes were identified in 86 (85.1 %) of the sequenced MDR-MRSA strains. Strains which harboured prophages had on average three prophages per genome (range 1-7) with a mean length of 25,151 bp. Only three strains (3.0 %) had prophages encoding an AMR gene (Fig 9), specifically the *dfrG* gene providing trimetoprim resistance. This gene was detected, in single or multiple copies, in the *Staphylococcus* phages ECel-2020q and phiSa2wa-st72 accordingly. The prophages identified were however associated with several known virulence factors. This included the well-known PVL toxin (encoded by *lukF-PV* and *lukS-PV*), δ-hemolysin (*hld)* and the enterotoxin-encoding genes *sea, sec2, sek, sel, sep* and *seq.* Prophages encoding the human immune evasion cluster were also detected, encoding the staphylokinase gene *(sak)* and the staphylococcal complement inhibitor (*scn)*. The prophages harbouring this cluster were phage 23MRA, ECel-2020g, P630, phiNM3, phiSAa119, SA1014ru, Sa2wa, SA345ru, SA7, SA780ru, SAP090B and tp310-3, which were present in 79 (78.2 %) of the sequenced MDR-MRSA strains.

**Fig 9.**
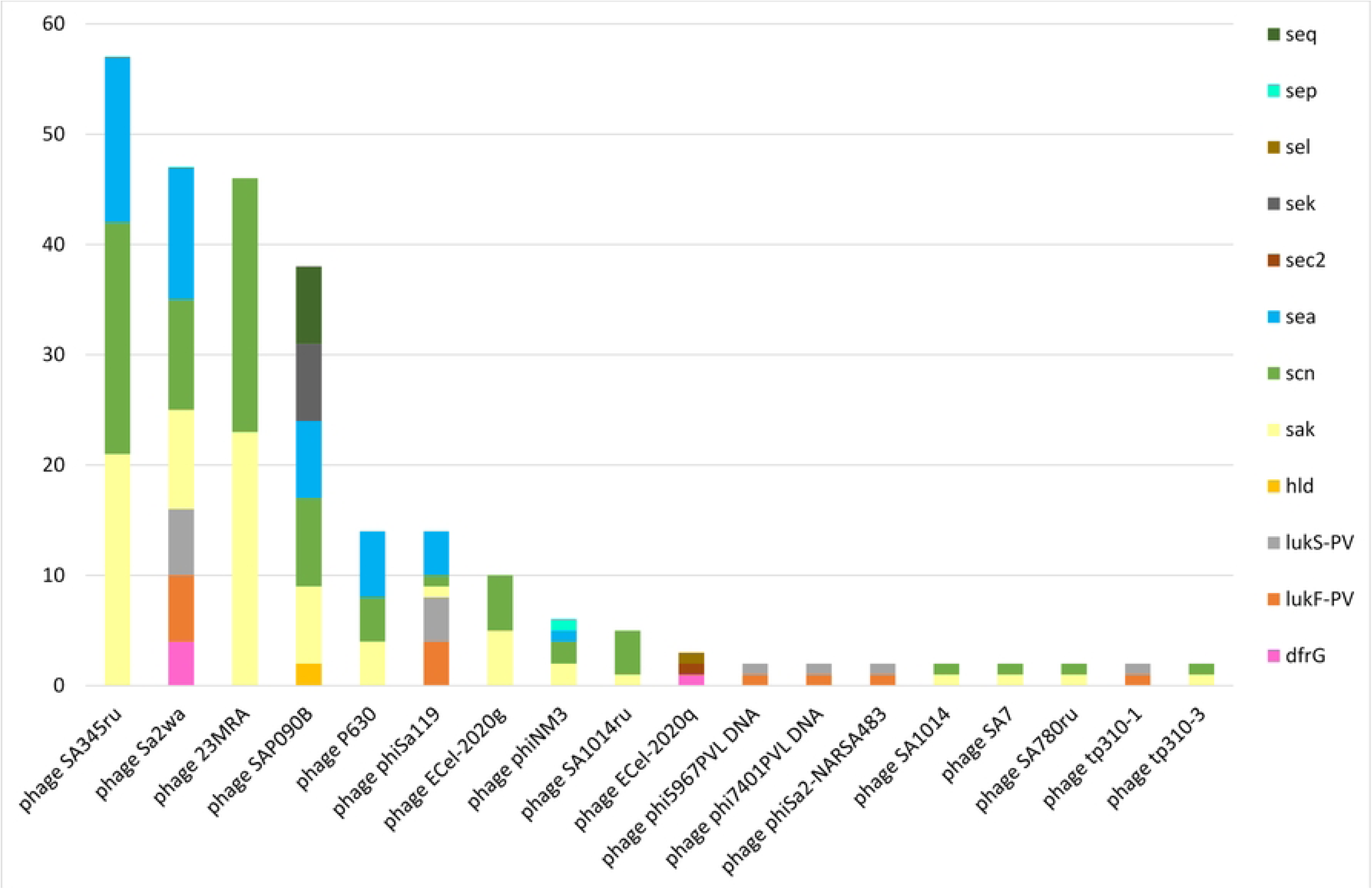
AMR and virulence genes in identified prophages in sequenced MDR-MRSA strains from Norway.

#### Composite transposons

A total of 207 composite transposons containing AMR-genes were identified in 76 (75.2 %) of the sequenced MDR-MRSA strains. Within these strains, we detected a median of one transposon per genome, with a mean length of 6,032 bp. All of these transposons contained AMR gene(s). The Tn*5405*, Tn*551* and Tn*552* were found in both the chromosome and on plasmids (Fig 10). The Tn*554*, Tn*558*, Tn*559*, Tn*6224* and Tn*917* were only found in the chromosomes of the analysed strains.

**Fig 10.**
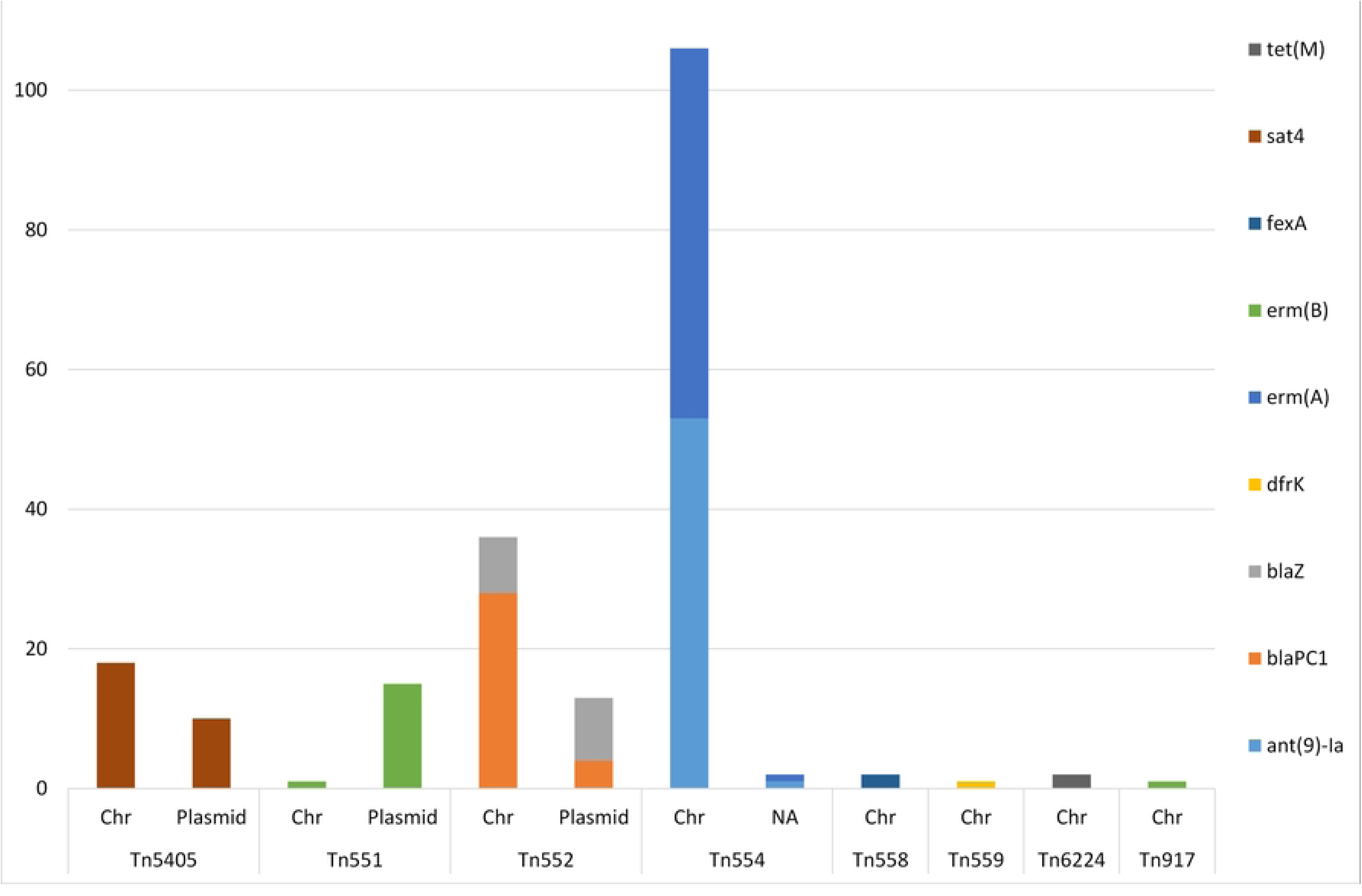
AMR genes identified in composite transposons in sequenced MDR-MRSA strains from Norway. Genomic location indicated by chromosome (Chr) or plasmid.

The AMR genes predominantly found within composite transposons were the aminoglycoside resistance gene *ant(9)-Ia* and the macrolide resistance gene *erm(A)* (n=54) (Fig 10). Additionally, the beta-lactam resistance genes *blaPC1* (n=32) and *blaZ* (n=17) were found in transposons, as well as the MLS gene *erm(B)* (n=17). Genes found more rarely in transposons were *tet(M)* (n=2) and *dfr(K)* (n=1).

Tn*554*-transposons (carrying the *ant(9)-Ia*- and *erm(A)* AMR genes), were present in one to four copies per strain. The Tn*551* and the Tn*552*-transposons was detected in a single or two copies when present in the strains. The Tn*664*-, Tn*5405*-, Tn*558*-, Tn*559* and Tn*917*-transposons, were present in a single copy per strain.

## Discussion

This study included all MDR-MRSA strains (n=452) detected in Norway during the study period 2008-2020, from a total of 23,412 MRSA strains. This project was made possible by the continued Norwegian MRSA surveillance effort that has been ongoing since 2008, with the aim of keeping multidrug-resistant pathogens from becoming endemic in healthcare institutions in Norway. Although the overall and yearly proportion of MDR-MRSA was relatively low, we observed an increase in the number of MDR-MRSA strains over time, as well as major changes in molecular epidemiology, during the study period. Specifically, we observed temporal shifts and predominace of certain genotypes (*spa*-types t1476, t127, t188 and t030/t037), indicating that these were successfully established and disseminating MDR-MRSA clones.

Although the information on place of acquisition for the MDR-MRSA strains is inherently limited by some uncertainty, it is interesting to note that only 13.1 % of strains were registered as aquired in Norway, while a majority were registered as aquired abroad. Compared to the overall MRSA population [12], this suggests import to be responsible for a comparatively larger proportion of MDR-MRSA strains, and similarly that Asia and Africa are the most frequent routes of transmission. This is also supported by the fact that we detect specific clones that have previously been reported as endemic or prevalent in these specific regions and countries [17, 18]. These findings highlight the importance of global cooperation, surveillance, antibiotic stewardship and infection control efforts that can limit both the emergence and global dissemination of these important pathogens.

A high proportion of the MDR-MRSA strains were healthcare-associated, mostly isolated from patients during hospital admission. The proportion of healthcare-associated strains was markedly higher among the MDR-MRSA strain collection than that reported in our surveillance study of the whole Norwegian MRSA population [12]. This suggests that the most highly resistant MRSA clones are more often hospital-associated than less resistant MRSA clones, although they have become more common in the community setting as well. This may among other factors be due to the high selection pressures and competition between MRSA clones provided by high antibiotic exposure [19]. MRSA are also endemic in many hospital environments wordwide [20], thus providing reservoirs for efficient spread into a vulnerable populations, especially in resource-limited settings with inadequate infection control [21]. A limitation to this study is the lack of temporal data on hospital and nursing home admissions, and thus that we had to rely on a broad definition of healthcare-associated MRSA. With this in mind, it is possible that the number of HA- and specifically hospital-associated cases are overestimated. However, the same definition of HA-MRSA was used in the surveillance study of the whole Norwegian MRSA population, in support of the relative differences observed in this study.

Interestingly, there was a larger proportion of MDR-MRSA strains from carriage than from clinical infections, compared to the corresponding numbers for the overall MRSA population in Norway [12]. This finding may reflect that a majority of these strains have been acquired abroad, and are thus (in Norway) mainly detected from asymptomatic carriers, e.g. due to screening upon contact with the healthcare system and differing screening practices. While it is possible that some MDR-MRSA strains have reduced virulence due to e.g. the physiological costs of maintaining resistance mechanisms, this can vary significantly between different strains and settings [22–24]. The prevalence of PVL-positive strains was 20.4 %, which was much lower than for the whole Norwegian MRSA strain population [12]. This also likely reflects the low number of infection compared to carriage strains, as PVL positive strains have typically been linked to clinical infections, whereas PVL-negative strains are more commonly associated with asymptomatic carriage [25].

In the MDR-MRSA strain collection, we observed almost universal resistance to antibiotics such as erythromycin and ciprofloxacin/norfloxacin in addition to cefoxitine. High levels of resistance were also observed for tetracycline, gentamicin, and clindamycin. Thus, in this group of strains, the choice of antibiotic treatment for potential infections is alarmingly and severely restricted. Resistance to mupirocin is however low, meaning that this is still an option for eradication/decolonization of a majority of these strains, while vancomycin and linezolid remain as treament options for all strains.

We observed a very large heteogeneity of *spa*-types and clonal complexes that were resistant towards five or six groups of antibiotics, indicating that multi-drug resistance is indeed a major challenge within the general MRSA population, not only in a few specific clones. The limited number of genotypes that were resistant to seven or eight groups of antibiotics however included well-known clones like MRSA CC239, that have been evolving antimicrobial resistance in high selection pressure environments for many decades, given their continous presence and global spread since the large hospital outbreaks of the 1970s [26, 27]. This underscores the critical importance of preventing the introduction of MRSA into hospitals and other healthcare facilities, especially in low-prevalence countries, and furthermore the importance of antibiotic stewardship and infection control in mitigating the risk of developing highly resistant strains within these environments.

Overall, we observed significant concordance between phenotypic (AST) and genotypic antimicrobial resistance profiles. In almost all cases, previously described AMR genes or mutations were detected, that could explain the observed phenotypes. Additionally, we noticed high clonality of resistance profiles, which corresponds well with the apparent clonality of specific MGEs carrying these AMR genes. Consequently, one can reasonably anticipate an antimicrobial profile based on genotyping results, although we do not suggest that antimicrobial susceptibility testing should be bypassed. On the other hand, certain genes and mutations which do not appear to confer phenotypic resistance in *S. aureus*, at least not to an extent which provides resistance according to clinical breakpoints, continue to be reported as resistance genes in the major AMR databases. Furthermore, especially for some groups of antibiotics, the genetics underlying the resistance phenotypes are still not well enough understood and/or are diffucult to interpret. This underscores the necessity for further investigation into the mechanisms underlying bacterial resistance as well as the importance of continuous curation and updating of AMR databases.

Nanopore sequencing technology was used to investigate the specific MGEs associated with AMR genes and mutations. Long-read sequencing techologies such as Oxford Nanopore Techologies (8) have facilitated a much more comprehensive investigation of MGEs, which can often be problematic to resolve by short-read sequencing (9). Consequently, our investigation revealed that AMR genes in MDR-MRSA strains are predominantly located within plasmids, as well as within staphylococcal cassette chromosome *mec* (SCC*mec*) elements and transposons, with a limited presence on prophages. The most common AMR genes were associated with wide range of MGEs, including different SCCmec types and multiple transposons and plasmids. Some MGEs were clonal, while others were widely distributed, indicating high potential for spread within the MRSA population. A majority of the plasmids were furthermore closely related, although differing in AMR traits, thus indiciating high plasticity. The most prevalent AMR phenotypes were furthermore caused by multiple AMR genes and/or mutations. As an example, we observed eight different genes or gene combinations in strains with MLS resistance. This large variation is likely a consequence of high selection pressure and thus multiple adaptations of MRSA clones over a long period of time, which now serves as an arsenal of genes available for providing resistance against different MLS antibiotics. This high diversity both in AMR genes and in acquisition mechanisms thus provide a significant advantage for dissemination of antibiotic resistance within the MRSA population. These findings highlight the complexity of resistance gene dissemination and the adaptive strategies employed by *S. aureus* in response to antimicrobial pressures.

## Materials and Methods

### Study design and population

This study is based on MRSA strains submitted to the Norwegian MRSA reference laboratory from 2008-2020. Inclusion was based on phenotypic resistance to five or more of the following antibiotic/resistance groups, with the antibiotic tested provided in parenteses; beta-lactams (cefoxitin), macrolide/lincosamide/streptogramin B (MLS) (erythromycin and clindamycin), aminoglycosides (gentamicin), tetracyclines (tetracycline), fucidanes (fusidic acid), fluoroquinolones (ciprofloxacin or norfloxacin), folate pathway inhibitors (trimethoprim-sulfamethoxazole, TMP-SMX), ansamycins (rifampicin), oxazolidinones (linezolid) and glycopeptides (vancomycin). Although all MRSA by current definitions [28] are regarded as multidrug-resistant (MDR), the strains included in this study are either not phenotypically resistant to, or have not been tested against, sufficient antibiotic groups to be designated as extensively drug-resistant (XDR). For lack of a better terminology, we thus refer to this strain collection as MDR-MRSA throughout this manuscript.

### Clinical and epidemiological data

Clinical and epidemiological data on all cases was collected from the Norwegian Surveillance System for Communicable Diseases (MSIS) and request forms from the referring laboratory or treating physician available from the laboratory information system (LIMS) at the national reference laboratory. The Information from MSIS included age, sex, admission to hospital or nursing home, place of acquisition and if it was part of a known outbreak (data from the Norwegian outbreak rapid alert system Vesuv) [9]. The data obtained from the LIMS included sample date, sampling site/type of sample and laboratory results. All MDR-MRSA cases were categorized as carriage, infection, invasive infection or unknown based on sampling site/type of sample. Age groups were defined according to Diaz et al. [10]. Due to lack of temporal data for hospitalized patients and nursing home stays, a case was classified as healthcare-associated (HA) if it was diagnosed during a hospital or nursing home stay, or if MDR-MRSA was detected in healthcare workersMLS (HCWs). Conversely, all other cases were classified as community-acquired (CA).

### Genotyping and antimicrobial susceptibility testing

Culturing, DNA extraction, *spa*-typing and assignment of *spa*-types to clonal complexes (CC) was performed as described previously [12]. The strains included in this study had previously undergone phenotypic antimicrobial susceptibility testing (AST) towards the following antibiotics: cefoxitin, erythromycin, clindamycin, fusidic acid, trimetoprim-sulfamethoxalate, tetracycline, gentamicin, ciprofloxacin/norfloxacin, rifampicin, mupirocin, linezolid and vancomycin. Susceptibility testing was performed as previously described [29] on all strains using the EUCAST (European Committee on Antimicrobial Susceptibility Testing) disk diffusion method and categorized as either susceptible, intermediate, or resistant according to the current EUCAST breakpoints at the time of testing. For clindamycin, inducible resistance was recorded as described in the EUCAST expert rules [30]. For vancomycin, the gradient strip test from BioMeriéux (2008-2013) or Liofilchem (2014-2020) was used.

### Whole genome sequencing

A selection of strains (n=101) were subjected to whole genome sequencing (WGS) by nanopore methodology, hereafter referred to as sequencing. These included all strains resistant to seven or more antibiotic groups (n=21), and a randomized selection among the remaining strains (n=80). Bacterial cells were first treated with proteinase K (2 mg/mL) and lysostaphin (0.1 mg/mL) for 15 min with shaking at 37 °C, before heating for 15 min at 65 °C. Genomic DNA was then isolated using the EZ1 DNA tissue kit with an EZ1 Advanced XL instrument (Qiagen). Sequencing libraries were prepared and multiplexed using the Rapid Barcoding Sequencing kit (SQK-RBK004), according to the RBK_9054_v2_revJ_14Aug2019 protocol. Sequencing libraries were loaded onto a R9.4.1 SpotON flow cell (FLO-MIN106D) and sequenced on a MinION Mk1B sequencer (Oxford Nanopore Technologies).

### Bioinformatic analyses

Dorado [31] v.0.4.2 was used for basecalling (SUP v3.6 model) and demultiplexing. Assembly was performed using Flye v.2.9.2 [32, 33]. Racon and Medaka (SUP v5.0.7 model) were used for polishing. Additionally, Homopolish [34] was used to remove possible systematic errors from the nanopore sequencing. The sequences are available from GenBank under BioProject ID PRJNA1186082.

Annotation was performed with Prokka v.1.14.6 [35]. Definition of the core genome and phylogeny were performed using Roary v3.13.0 with default settings [36] and FastTree with the GTR model [37].

Plasmid classification was performed by MOB-suite v.3.1.8 [38], with the typing and clustering modules. Mashtree [39] was then used to construct a plasmid phylogeny. Plasmids were furthermore merged into communities/subcommunities using the Pling bioinformatic tool [40].

Putative prophages were detected using Phastest [41] including only intact phages, and identified using nBLAST against reference *S. aureus* phages in the GenomeNet Virus-Host DataBase (VHDB) [42]. Transposons and IS-elements were detected using the MobileElementFinder [43]. *SCCmec* chromosome cassettes were typed using SCCmecFinder [44]. Whole *SCCmec* elements were extracted by *in silico* PCR with SeqKit [45], using modified primers from Ito *et al* [46].

For whole genomes as well as for specific MGEs, AMR genes and point mutations were detected using AMRFinderPlus v3.10.30 with organism-specific settings for *S. aureus*, and cut-off of 90 % on protein identity and 50 % on coverage [47].

### Visualization

Upset plots were created using R studio v4.3.3 with the ggplot2 [48], ComplexUpset [49, 50] and cowplot [51] packages. Visualization of the core genome tree with phenotypic and genotypic traits was performed with iTol v.5 [52]. The world map was created using GeoPandas [53] and Matplotlib [54].

## Ethical statement

The project was approved, with exemption from informed consent of participants, by the Norwegian Regional Committees for Medical and Health Research Ethics (REC) South-East with reference number 352380. A Data Protection Impact Assessment was performed and approved by the responsible institution, the Clinic of Laboratory Medicine at St. Olavs Hospital, Trondheim University Hospital.

## Acknowledgements

The authors would like to acknowledge the contribution of all submitting medical microbiology laboratories to the continued MRSA surveillance effort in Norway, as well as the laboratory personnel who have contributed to the analysis of these strains at the Norwegian MRSA reference laboratory at St. Olavs Hospital.

## References

1. Howden BP, Giulieri SG, Wong Fok Lung T, Baines SL, Sharkey LK, Lee JYH, et al. Staphylococcus aureus host interactions and adaptation. Nat Rev Microbiol. 2023;21(6):380–95.

2. Boucher HW, Corey GR. Epidemiology of methicillin-resistant Staphylococcus aureus. Clin Infect Dis. 2008;46 Suppl 5:S344–9.

3. Richardson EJ, Bacigalupe R, Harrison EM, Weinert LA, Lycett S, Vrieling M, et al. Gene exchange drives the ecological success of a multi-host bacterial pathogen. Nat Ecol Evol. 2018;2(9):1468–78.

4. Zhao W, Zeng W, Pang B, Luo M, Peng Y, Xu J, et al. Oxford nanopore long-read sequencing enables the generation of complete bacterial and plasmid genomes without short-read sequencing. Frontiers in Microbiology. 2023;14.

5. Mores CR, Montelongo C, Putonti C, Wolfe AJ, Abouelfetouh A. Investigation of Plasmids Among Clinical Staphylococcus aureus and Staphylococcus haemolyticus Isolates From Egypt. Front Microbiol. 2021;12:659116.

6. Ito T, Kuwahara-Arai K, Katayama Y, Uehara Y, Han X, Kondo Y, et al. Staphylococcal Cassette Chromosome mec (SCCmec) analysis of MRSA. Methods Mol Biol. 2014;1085:131–48.

7. Katayama Y, Ito T, Hiramatsu K. Genetic organization of the chromosome region surrounding mecA in clinical staphylococcal strains: role of IS431-mediated mecI deletion in expression of resistance in mecA-carrying, low-level methicillin-resistant Staphylococcus haemolyticus. Antimicrob Agents Chemother. 2001;45(7):1955–63.

8. Ito T, Katayama Y, Hiramatsu K. Cloning and nucleotide sequence determination of the entire mec DNA of pre-methicillin-resistant Staphylococcus aureus N315. Antimicrob Agents Chemother. 1999;43(6):1449–58.

9. Leinweber H, Sieber RN, Larsen J, Stegger M, Ingmer H. Staphylococcal Phages Adapt to New Hosts by Extensive Attachment Site Variability. mBio. 2021;12(6):e0225921.

10. Aslam B, Khurshid M, Arshad MI, Muzammil S, Rasool M, Yasmeen N, et al. Antibiotic Resistance: One Health One World Outlook. Front Cell Infect Microbiol. 2021;11:771510.

11. Murray CJL, Ikuta KS, Sharara F, Swetschinski L, Robles Aguilar G, Gray A, et al. Global burden of bacterial antimicrobial resistance in 2019: a systematic analysis. The Lancet. 2022;399(10325):629–55.

12. Ronning TG, Enger H, Afset JE, As CG. Insights from a decade of surveillance: Molecular epidemiology of methicillin-resistant Staphylococcus aureus in Norway from 2008 to 2017. PLoS One. 2024;19(3):e0297333.

13. Guerillot R, Goncalves da Silva A, Monk I, Giulieri S, Tomita T, Alison E, et al. Convergent Evolution Driven by Rifampin Exacerbates the Global Burden of Drug-Resistant Staphylococcus aureus. mSphere. 2018;3(1).

14. Howe RA, Wootton M, Noel AR, Bowker KE, Walsh TR, MacGowan AP. Activity of AZD2563, a novel oxazolidinone, against Staphylococcus aureus strains with reduced susceptibility to vancomycin or linezolid. Antimicrob Agents Chemother. 2003;47(11):3651–2.

15. Liakopoulos A, Spiliopoulou I, Damani A, Kanellopoulou M, Schoina S, Papafragas E, et al. Dissemination of two international linezolid-resistant Staphylococcus epidermidis clones in Greek hospitals. J Antimicrob Chemother. 2010;65(5):1070–1.

16. Johnson J, Soehnlen M, Blankenship HM. Long read genome assemblers struggle with small plasmids. Microb Genom. 2023;9(5).

17. Lawal OU, Ayobami O, Abouelfetouh A, Mourabit N, Kaba M, Egyir B, et al. A 6-Year Update on the Diversity of Methicillin-Resistant Staphylococcus aureus Clones in Africa: A Systematic Review. Front Microbiol. 2022;13:860436.

18. Mohamad Farook NA, Argimon S, Abdul Samat MN, Salleh SA, Sulaiman S, Tan TL, et al. Diversity and Dissemination of Methicillin-Resistant Staphylococcus aureus (MRSA) Genotypes in Southeast Asia. Trop Med Infect Dis. 2022;7(12).

19. de Vos AS, de Vlas SJ, Lindsay JA, Kretzschmar MEE, Knight GM. Understanding MRSA clonal competition within a UK hospital; the possible importance of density dependence. Epidemics. 2021;37:100511.

20. Stefani S, Chung DR, Lindsay JA, Friedrich AW, Kearns AM, Westh H, et al. Meticillin-resistant Staphylococcus aureus (MRSA): global epidemiology and harmonisation of typing methods. International Journal of Antimicrobial Agents. 2012;39(4):273–82.

21. Christopher S, Verghis RM, Antonisamy B, Sowmyanarayanan TV, Brahmadathan KN, Kang G, et al. Transmission dynamics of methicillin-resistant Staphylococcus aureus in a medical intensive care unit in India. PLoS One. 2011;6(7):e20604.

22. Rudkin JK, Edwards AM, Bowden MG, Brown EL, Pozzi C, Waters EM, et al. Methicillin Resistance Reduces the Virulence of Healthcare-Associated Methicillin-Resistant Staphylococcus aureus by Interfering With the agr Quorum Sensing System. The Journal of Infectious Diseases. 2012;205(5):798–806.

23. Taglialegna A, Varela MC, Rosato RR, Rosato AE. VraSR and Virulence Trait Modulation during Daptomycin Resistance in Methicillin-Resistant *Staphylococcus aureus* Infection. mSphere. 2019;4(1):10.1128/msphere.00557-18.

24. Rao Y, Peng H, Shang W, Hu Z, Yang Y, Tan L, et al. A vancomycin resistance-associated WalK(S221P) mutation attenuates the virulence of vancomycin-intermediate Staphylococcus aureus. J Adv Res. 2022;40:167–78.

25. Boyle-Vavra S, Daum RS. Community-acquired methicillin-resistant Staphylococcus aureus: the role of Panton–Valentine leukocidin. Laboratory Investigation. 2007;87(1):3–9.

26. Shanson DC. Antibiotic-resistant Staphylococcus aureus. Journal of Hospital Infection. 1981;2:11–36.

27. Monecke S, Slickers P, Gawlik D, Müller E, Reissig A, Ruppelt-Lorz A, et al. Molecular Typing of ST239-MRSA-III From Diverse Geographic Locations and the Evolution of the SCCmec III Element During Its Intercontinental Spread. Frontiers in Microbiology. 2018;9.

28. Magiorakos AP, Srinivasan A, Carey RB, Carmeli Y, Falagas ME, Giske CG, et al. Multidrug-resistant, extensively drug-resistant and pandrug-resistant bacteria: an international expert proposal for interim standard definitions for acquired resistance. Clinical Microbiology and Infection. 2012;18(3):268–81.

29. Enger H, Larssen KW, Damås ES, Aamot HV, Blomfeldt A, Elstrøm P, et al. A tale of two STs: molecular and clinical epidemiology of MRSA t304 in Norway 2008-2016. Eur J Clin Microbiol Infect Dis. 2022;41(2):209–18.

30. EUCAST. The European Committee on Antimicrobial Susceptibility Testing Expert Rules v 3.2 http://www.eucast.org/clinical_breakpoints/. 2023.

31. Samarakoon H, Ferguson JM, Gamaarachchi H, Deveson IW. Accelerated nanopore basecalling with SLOW5 data format. Bioinformatics. 2023;39(6).

32. Kolmogorov M, Yuan J, Lin Y, Pevzner PA. Assembly of long, error-prone reads using repeat graphs. Nat Biotechnol. 2019;37(5):540–6.

33. Lin Y, Yuan J, Kolmogorov M, Shen MW, Chaisson M, Pevzner PA. Assembly of long error-prone reads using de Bruijn graphs. Proc Natl Acad Sci U S A. 2016;113(52):E8396–E405.

34. Huang YT, Liu PY, Shih PW. Homopolish: a method for the removal of systematic errors in nanopore sequencing by homologous polishing. Genome Biol. 2021;22(1):95.

35. Seemann T. Prokka: rapid prokaryotic genome annotation. Bioinformatics. 2014;30(14):2068–9.

36. Page AJ, Cummins CA, Hunt M, Wong VK, Reuter S, Holden MT, et al. Roary: rapid large-scale prokaryote pan genome analysis. Bioinformatics. 2015;31(22):3691–3.

37. Price MN, Dehal PS, Arkin AP. FastTree: Computing Large Minimum Evolution Trees with Profiles instead of a Distance Matrix. Molecular Biology and Evolution. 2009;26(7):1641–50.

38. Robertson J, Nash JHE. MOB-suite: software tools for clustering, reconstruction and typing of plasmids from draft assemblies. Microb Genom. 2018;4(8).

39. Katz LS, Griswold T, Morrison SS, Caravas JA, Zhang S, den Bakker HC, et al. Mashtree: a rapid comparison of whole genome sequence files. J Open Source Softw. 2019;4(44).

40. Frolova D, Lima L, Roberts L, Bohnenkämper L, Wittler R, Stoye J, et al. Applying rearrangement distances to enable plasmid epidemiology with pling. bioRxiv. 2024:2024.06.12.598623.

41. Wishart DS, Han S, Saha S, Oler E, Peters H, Grant Jason R, et al. PHASTEST: faster than PHASTER, better than PHAST. Nucleic Acids Research. 2023;51(W1):W443–W50.

42. Mihara T, Nishimura Y, Shimizu Y, Nishiyama H, Yoshikawa G, Uehara H, et al. Linking Virus Genomes with Host Taxonomy. Viruses. 2016;8(3):66.

43. Johansson MHK, Bortolaia V, Tansirichaiya S, Aarestrup FM, Roberts AP, Petersen TN. Detection of mobile genetic elements associated with antibiotic resistance in Salmonella enterica using a newly developed web tool: MobileElementFinder. Journal of Antimicrobial Chemotherapy. 2020;76(1):101–9.

44. Kaya H, Hasman H, Larsen J, Stegger M, Johannesen TB, Allesøe RL, et al. SCC *mec*Finder, a Web-Based Tool for Typing of Staphylococcal Cassette Chromosome *mec* in Staphylococcus aureus Using Whole-Genome Sequence Data. mSphere. 2018;3(1):10.1128/msphere.00612-17.

45. Shen W, Le S, Li Y, Hu F. SeqKit: A Cross-Platform and Ultrafast Toolkit for FASTA/Q File Manipulation. PLOS ONE. 2016;11(10):e0163962.

46. Ito T, Katayama Y, Asada K, Mori N, Tsutsumimoto K, Tiensasitorn C, et al. Structural comparison of three types of staphylococcal cassette chromosome mec integrated in the chromosome in methicillin-resistant Staphylococcus aureus. Antimicrob Agents Chemother. 2001;45(5):1323–36.

47. Feldgarden M, Brover V, Gonzalez-Escalona N, Frye JG, Haendiges J, Haft DH, et al. AMRFinderPlus and the Reference Gene Catalog facilitate examination of the genomic links among antimicrobial resistance, stress response, and virulence. Sci Rep. 2021;11(1):12728.

48. Wickham H. ggplot2: Elegant Graphics for Data Analysis: Springer-Verlag New York; 2016.

49. Krassowski M. ComplexUpset. https://doiorg/105281/zenodo3700590. 2020.

50. Lex A, Gehlenborg N, Strobelt H. UpSet: Visualization of Intersecting Sets,. IEEE Transactions on Visualization and Computer Graphics. 2014;20(12):1983–92.

51. Wilke CO. cowplot: Streamlined Plot Theme and Plot Annotations for ‘ggplot2’. R package version 1.1.3. 2024.

52. Letunic I, Bork P. Interactive Tree Of Life (iTOL) v5: an online tool for phylogenetic tree display and annotation. Nucleic Acids Research. 2021;49(W1):W293–W6.

53. Jordahl K, Bossche JVd, Fleischmann M, Wasserman J, McBride J, Tratner JG, et al. geopandas/geopandas: v0.8.1. Zenodo. 2020.

54. Hunter JD. Matplotlib: A 2D Graphics Environment. Computing in Science & Engineering. 2007;vol. 9, no. 3, pp. 90–95, May-June 2007.

